# A Rab-Kinesin12-Fused kinase module couples vesicle delivery and phragmoplast remodelling during cytokinesis in *Arabidopsis*

**DOI:** 10.1101/2024.10.11.617818

**Authors:** Liam Elliott, Monika Kalde, Moé Yamada, Michal Hála, Andrei Smertenko, Frédérique Rozier, James Evry, Niloufer Irani, Yvon Jaillais, Patrick J Hussey, Viktor Žárský, Ian Moore, Charlotte Kirchhelle

## Abstract

Cytokinesis is a key process in the development of multicellular organisms, through both formative and proliferative divisions. In land plants, successful cytokinesis requires precise targeting of Golgi-derived vesicles to the future division plane by the phragmoplast, a cytoskeletal structure that undergoes continuous remodelling. Endomembrane trafficking and phragmoplast remodelling are tightly coupled, indicating the existence of active crosstalk. However, although many molecular regulators of membrane trafficking and cytoskeletal organisation are known, it remains poorly understood how these processes are co-ordinated. Here, we describe a regulatory module consisting of the membrane-associated GTPase RAB-A2a, cytoskeleton-associated Class II Kinesin-12s, and the Fused kinase orthologue TIO. We provide evidence that the interaction between these molecules at the midzone is essential for cytokinesis in *Arabidopsis* through simultaneously targeting vesicles to the midzone and coupling phragmoplast remodelling to vesicle delivery.

## Introduction

During cytokinesis in plant cells, both the endomembrane and cytoskeleton systems undergo substantial remodelling, which differs markedly from that found in other eukaryotic lineages. Following the dissolution of the spindle at the end of anaphase, plant cells form a phragmoplast, a cytoskeletal structure unique to land plants and some green algae. The phragmoplast is predominantly composed of microtubules which are arranged into two partially-overlapping arrays, with microtubule plus-ends directed towards the future division site at the midzone. Endomembrane vesicles from the *trans*-Golgi network/early endosome (TGN/EE) are mass-transported along phragmoplast microtubules to the midzone where they fuse to form the cell plate, an endomembrane compartment also unique to the green lineage that is filled with oligosaccharides. The phragmoplast initially forms a disk, but subsequently undergoes centrifugal expansion to form a ring. Phragmoplast expansion is achieved through microtubule depolymerisation at the phragmoplast core once the cell plate no longer requires a cytoskeletal scaffold, and simultaneous microtubule polymerisation at the phragmoplast outer edge. This remodelling drives simultaneous centrifugal expansion of the cell plate, which eventually fuses with the plasma membrane to form a new crosswall^1^.

To ensure successful cytokinesis, the cell plate needs to expand within a 2D plane to form an organelle large enough to partition the entire cell (new cross walls are typically several hundred µm^2^). This process depends on the precise control of two factors: (1) spatial targeting of vesicles to the midzone via the phragmoplast, and (2) temporal coordination between membrane delivery and phragmoplast remodelling. Cytoskeleton-associated molecular machinery located at the phragmoplast and cell periphery regulate phragmoplast formation, remodelling and positioning. These include kinesins of the Kinesin-12 family^2,3^, IQ67 domain family proteins^4^, microtubule crosslinkers of the MICROTUBULE-ASSOCIATED PROTEIN 65 (MAP65) family^5–8^, and diverse regulators of microtubule nucleation, polymerisation and severing (summarized by Smertenko and colleagues^9^). Microtubule depolymerisation at the phragmoplast core is regulated by at least three signalling pathways involving Mitogen-Activated Protein Kinase (MAPK), Cyclin-dependent kinase (CDK), and Aurora Kinase^10–15^. The widely-conserved signalling kinase TWO-IN-ONE (TIO) has also been linked to initiation of phragmoplast expansion in *Arabidopsis* microspores^16^, a function that may relate to its interactions with phragmoplast-associated kinesins PAKRP1/Kinesin-12A (Kin-12A) and PAKRP1L/Kinesin-12B (Kin-12B), and TETRASPORE^17,18^. Kin-12A/-12B localise to interdigitating microtubules at the phragmoplast midzone in somatic cells of *Arabidopsis thaliana*^19,20^, and are redundantly required for proper phragmoplast formation during mitotic cytokinesis of *Arabidopsis* microspores^3^. In a *kin12a kin12b* (*kin12a/b*) double null mutant, the majority of *Arabidopsis* microspores fail to form organised phragmoplasts during cytokinesis, causing reduced fertility^3^. However, somatic cytokinesis has been reported to occur normally in *kin12a/b* double mutants, indicating cell-type specific functional redundancy^3^.

Directional transport of TGN/EE membrane vesicles along phragmoplast microtubules to the midzone is believed to be driven by phragmoplast-associated transport kinesins. Such kinesins have not been identified in angiosperms, but recent data indicate that in the moss *Physcomitirum patens*, these include members of the class II Kin-12 family^21^. Vesicle transport to the cell plate also requires regulators of endomembrane trafficking such as Rab GTPases, multi-subunit tethering complexes, dynamins, and SNARE components^22–30^. Vesicles arrive at the phragmoplast midzone at the leading edge, before fusing to form a tubulo-vesicular network in the transition zone, and eventually becoming complete cell plate in the core phragmoplast lagging zone. While phragmoplast expansion is essential for patterning vesicle transport during cell plate formation, vesicle trafficking from the TGN/EE is also conversely required for phragmoplast expansion. Pharmacological inhibition of membrane trafficking from the TGN/EE^31^ as well as mutants in endomembrane regulators^32,33^ can perturb phragmoplast organisation and expansion, indicating the existence of crosstalk between mechanisms controlling cytoskeleton remodelling and endomembrane trafficking during cytokinesis to ensure co-ordination of both processes. However, it is not known how information about endomembrane dynamics and activity during cytokinesis is integrated into control of phragmoplast formation and dynamics.

Here, we describe the interaction between the endomembrane-associated small GTPase RAB-A2a, members of the cytoskeleton-associated Kinesin-12 family, and the Fused kinase orthologue TIO during cytokinesis in plant cells. We present evidence that their interaction forms a regulatory module that functions to both precisely target molecular regulators of cytokinesis to the phragmoplast midzone, as well as coupling cytoskeletal remodelling to endomembrane delivery during cell division. We conclude the RAB-A2a-Kin-12-TIO module provides a mechanistic basis for robust spatio-temporal control of plant cytokinesis.

## Results

### RAB-A2a interacts with Class II Kinesin-12s *in vitro* and *in planta*

In a Y2H screen for interactors of RAB-A2a, we isolated 11 independent clones encoding for the C-terminal/tail regions of three Kinesin-12 proteins: Kin-12A, Kin-12B and Kin-12F (Figure 1A). The six Kinesin-12 proteins (Kin-12s) encoded in the *Arabidopsis* genome are grouped into two subclasses, Class I and Class II^34^. In our Y2H screen, we identified all three members of Class II, while no clones corresponding to Class I Kin-12s or any other kinesins were isolated. To validate the interaction between RAB-A2a and Class II Kin-12s, we conducted pairwise Y2H tests using independently-cloned constructs of the Kin-12 tail regions, and confirmed interactions for all Class II Kin-12s with wild-type RAB-A2a as well as RAB-A2a[QL], a mutant variant with reduced GTPase activity that is considered ‘constitutively active’^25^ (Figure 1AB, S1A). By contrast, we found no evidence of a Y2H interaction between Class II Kin-12 tail regions and RAB-A2a[SN], a variant predicted to have increased binding affinity for GDP and thus be arrested upstream of Rab GTPase activation^25^ (Figure 1B, S1A). We also found reduced interaction with RAB-A2a[NI], a variant with reduced nucleotide affinity^25^ (Figure 1B, S1A). We also tested interactions between the Class II Kin-12 tail regions and other Rab GTPases that localize to the cell plate during cytokinesis, and found no interaction between Kin-12A, -12B, or 12F and wild-type or [QL] variants of RAB-A5c, RAB-A3, and RAB-E1d (Figure 1B, S1A). Taken together, these data indicate that Class II Kin-12s can interact with RAB-A2a in its active form.

**Figure 1:**
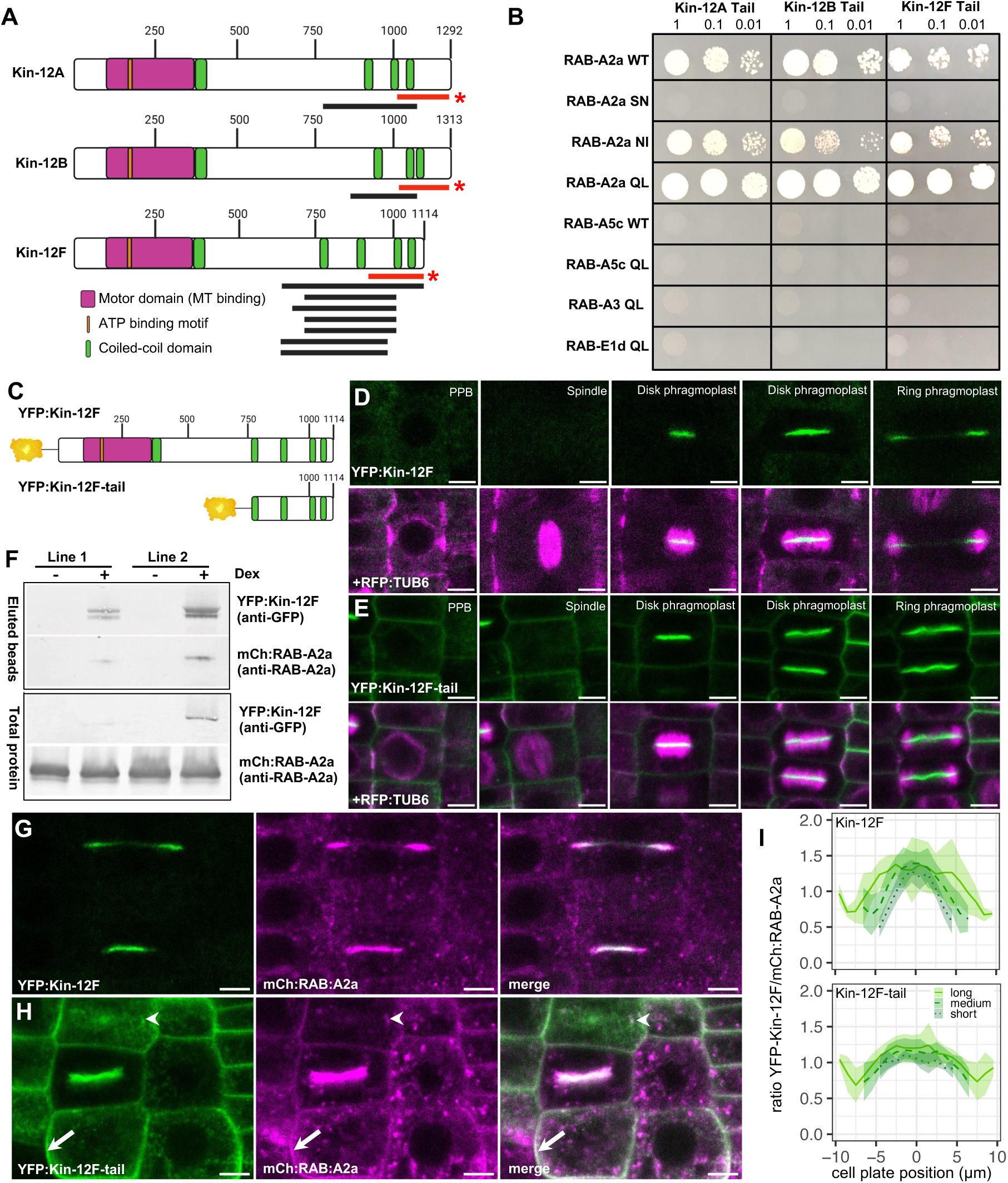
RAB-A2a interacts with lass II kinesin-12 members *in vitro* and *in planta*. **(A)** Schematic depiction of Class II Kin-12 proteins. Black lines: clone region isolated as interactors of RAB-A2a in initial Y2H screen. Red lines and astericks: independently cloned regions used in pairwise Y2H tests in (B). Schematic depictions inspired by those in^32^ **(B)** Pairwise Y2H tests between Kin-12 tail regions and Rab-A GTPase variants on SD-Leu-Trp-Ade-His. **(C)** Schematic depiction of YFP:Kin-12F and YFP:Kin-12F-tail. **(D,E)** Confocal Laser Scanning Microscopy (CLSM) sections of primary root meristematic epidermal cells co-expressing RFP:TUB6 and DEX>>YFP:Kin-12F (D) or DEX>>YFP:Kin-12F-tail (E) at different stages of cell division. **(F)** Immunoblots analysed with anti-YFP and anti-RAB-A2a showing co-immunoprecipitation between YFP:Kin-12F and mCh:RAB-A2a in two independent lines co-expressing DEX>>YFP:Kin-12F and mCh:RAB-A2a, in presence (+) or absence (-) of Dex to induce expression of YFP:Kin-12F. **(G,H)** CLSM section of primary root meristematic epidermal cells co-expressing *mCh:RAB-A2a* and *DEX>>YFP:Kin-12F* (G) or *DEX>>YFP:Kin-12F-tail* (H). White arrow indicates PM, arrowhead indicates intracellular compartment. **(I)** Fluorescence intensity ratio between YFP:Kin-12F (top) or YFP:Kin-12F-tail (bottom) and mCh:RAB-A2a along cell plates as those shown in (G,H). Cell plates were grouped by diameter into short (<9µm), medium (9-13µm) and long (>13µm). Lines are mean values, shaded areas are +/- 1SD. N = 4 (YFP:Kin-12F-tail short), 7 (YFP:Kin-12F long), 10 (YFP:Kin-12F short), 11 (YFP:Kin-12F-tail long), 14 (YFP:Kin-12F medium), 16 (YFP:Kin-12F-tail medium). Scale bars, 5µm. Graphics created with biorender.com.

Kin-12A and Kin-12B localize to the phragmoplast midzone during cytokinesis, and are thought to play a structural role in phragmoplast organization in microspores^3,19,20^. To investigate the role of the related Kin-12F, we generated stable transgenic lines expressing fluorescently-tagged Kin-12F. We first attempted to express fluorescently-tagged Kin-12F constitutively under the control of the 35S promoter. However, we recovered no stable transgenic lines with either a gDNA or cDNA amplification template. We therefore used the dexamethasone-inducible pOp6/LhGR expression system driven by the *AtRPS5a* promoter^35^ to conditionally express YFP:Kin-12F (*AtRPS5a>>Dex>>YFP:Kin-12F*), as well as a truncated protein variant encompassing the Kin-12F tail including the RAB-A2a interaction domain identified in Y2H, but lacking the Kin-12F motor domain (*AtRPS5a>>Dex>>YFP:Kin-12F-tail*; Figure 1C). We observed YFP fluorescence in 6 of 18 *AtRPS5a>>Dex>>YFP:Kin-12F* T2 lines after Dex treatment, and 6 of 8 *AtRPS5a>>Dex>>YFP:Kin-12F-tail* T2 lines after Dex treatment.

After 24h induction, YFP:Kin-12F localised to the phragmoplast midzone during cytokinesis in primary roots, where it closely followed the expanding phragmoplast labelled with RFP:TUB-6 (Figure 1D), matching previous reports of Kin-12A and -12B localisation^3,20^. We did not detect YFP:Kin-12F during earlier stages of cell division when preprophase bands and spindles were present, or in interphase cells (Figure 1D, S1C), although the AtRPS5a-driven pOp6/LhGR was active during all stages of the cell cycle in meristematic root cells (Figure S1B). By contrast, we were able to detect YFP:Kin-12F-tail during all stages of the cell cycle (Figure 1E, S1D), suggesting that (1) Kin-12F is post-transcriptionally or - translationally suppressed in non-dividing cells and (2) this suppression requires the N-terminal end of Kin-12F. Unlike the full-length YFP:Kin-12F, YFP:Kin-12F-tail localised to the plasma membrane (PM) in interphase, and labelled the entire cell plate during cytokinesis (Figure 1E, S1D). To confirm the localisation pattern of Kin-12F that we observed when expressed from the *AtRPS5a* promoter, we also performed immunolocalisation of endogenous Kin-12F in wild-type *Arabidopsis* primary roots using a polyclonal antibody against a 16 amino acid peptide sequence located in the N-terminal region of Kin-12F. During cytokinesis, this anti-Kin-12F antibody was detected at the phragmoplast midzone of both disk- and ring-phase phragmoplasts, similar to the localisation pattern of conditionally expressed YFP:Kin-12F (Figure 1D, S1G,H). This signal was abolished in primary roots of a *kin-12f* SALK mutant, indicating the antibody was specific to Kin-12F (Figure S1G).

We next co-expressed *AtRPS5a>>Dex>>YFP:Kin-12F/YFP:Kin-12F-tail* with *p35S::mCherry:RAB-A2a*, and performed co-immunoprecipitation experiments against YFP:Kin-12F/YFP:Kin-12F-tail. In these experiments, we found that mCh:RAB-A2a co-purified with YFP:Kin12F (Figure 1F), as well as YFP:Kin-12F-tail (Figure S1E), demonstrating these proteins can interact *in planta*.

### Class II Kin-12 members and RAB-A2a can mutually influence each other’s patterning

To investigate the functional relevance of these interactions, we performed quantitative co-localisation analyses between mCh:RAB-A2a and YFP:Kin-12F/YFP:Kin-12F-tail at the cell plate. In the presence of YFP:Kin-12F, mCh:RAB-A2a labelled the entirety of early cell plates, and was later enriched at the edges of expanding cell plates (Figure 1G, S2A). YFP:Kin-12F broadly followed the same pattern, but was depleted relative to mCh:RAB-A2a at the leading edge, which is the major site of vesicle delivery (Figure 1G, I, S2A). However, when co-expressed with YFP:Kin-12F-tail, mCh:RAB-A2a distribution was notably altered towards a more uniform pattern across the cell plate (Figure S2B). This closely matched the pattern of YFP:Kin-12F-tail at cell plates (Figure 1E,H, I). Furthermore, both YFP:Kin-12F-tail and mCh:RAB-A2a colocalised in interphase cells at the PM, as well as occasionally at intracellular punctae (Figure 1H, white arrowheads/arrows, S1D, S2D). We did not observe substantial PM localisation for mCh:RAB-A2a in the absence of YFP:Kin-12F-tail (Figure S1C, S2C), indicating that this localisation shift was driven by the presence of YFP:Kin-12F-tail. PM localisation of YFP:Kin-12F-tail and mCh:RAB-A2a were both abolished by treatment with the TGN/EE recycling inhibitor Brefeldin-A (Figure S2E,F) as well as by expression of the dominant-negative protein variants RAB-A2a[NI] and RAB-A2a[SN] (Figure S2G-L), indicating that PM localisation of YFP:Kin-12F-tail was dependent on RAB-A2a-mediated exocytic transport from the TGN/EE^25,36^. Taken together, our data demonstrates that Class II Kin-12 members and RAB-A2a interact *in planta* at the cell plate, and can mutually influence each other’s localisation.

### Class II Kinesin-12s are redundantly required for somatic cytokinesis

Kin-12A and -12B are required for phragmoplast and cell plate formation in *Arabidopsis* microspores^3^. However, *kin12a/b* mutants have been reported to have no visible defects in somatic cytokinesis, indicating that they may act redundantly with other kinesins during cytokinesis in somatic cells^3,37^. To test this hypothesis, we used CRISPR-Cas9 to abolish Kin-12F function in the *kin12a/b* background^3^. To avoid possible lethality, we targeted Kin-12F via CRIPSR-Cas9 driven from the *GLABRA2 (GL2)* promoter, whose activity in roots is confined to atrichoblast cell files^38^ (*pGL2::kin-12f:cas9-nls-mturquoise*). To confirm suppression of Kin-12F by *pGL2::kin-12f::cas9-nls-mturquoise*, we first introduced it into the YFP:Kin-12F background. The cas9-nls-mturquoise signal was detected exclusively in atrichoblast cell files as expected, where it was present in the nuclei during interphase and dispersed throughout the cytoplasm during mitosis following dissolution of the nucleus, before gradually returning to the re-forming nuclei during telophase (Figure 2A,C-E, Figure S5).

**Figure 2:**
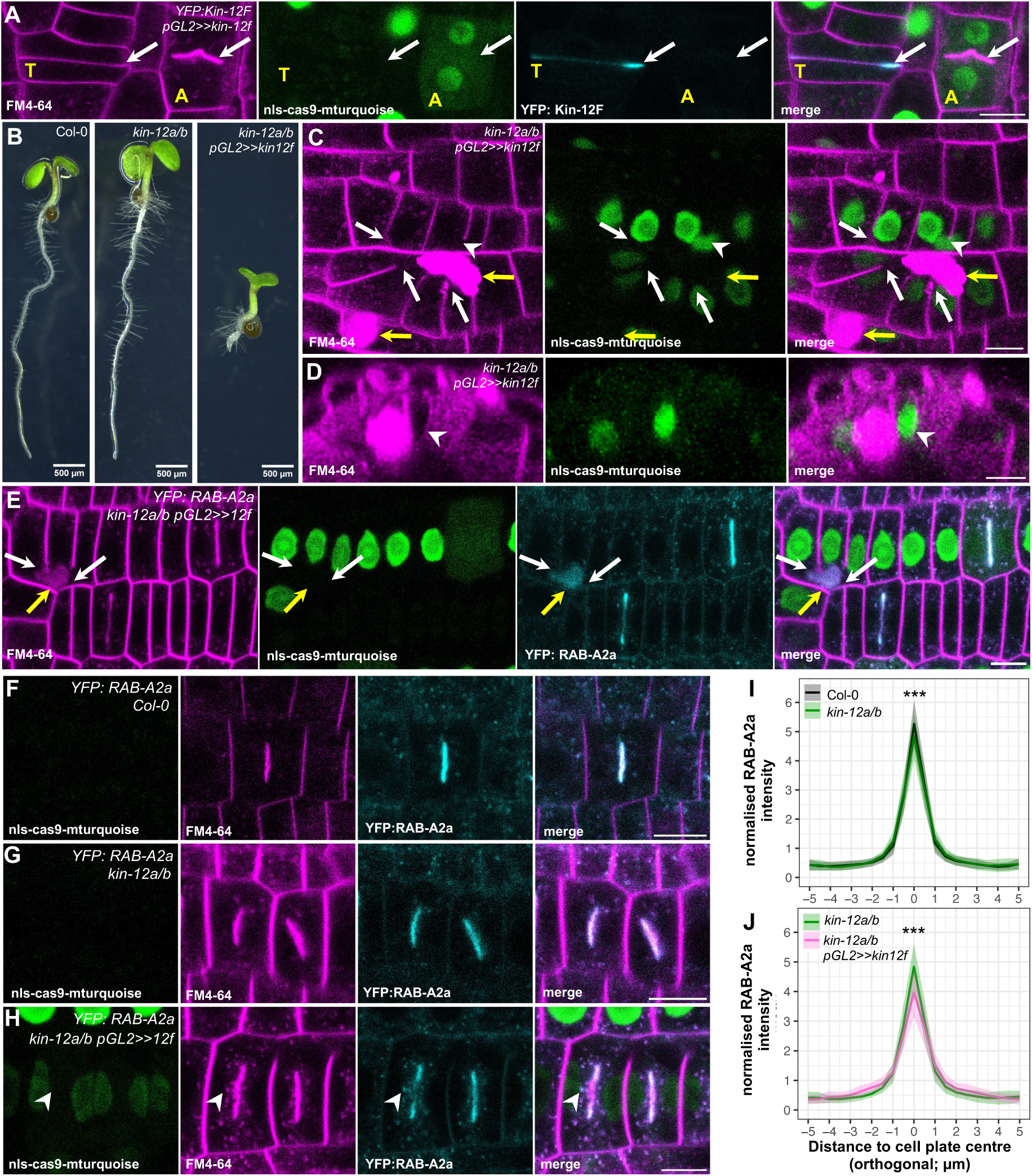
Class II Kin-12 proteins are required for somatic cytokinesis and contribute to midzone membrane targeting. **(A)** CLSM section of primary root meristematic epidermal cells in DEX>>YFP:Kin-12F*, pGL2::kin-12f::cas9-nls-mturquoise (pGL2>>12f)* background counterstained with FM4-64. White arrows indicate cell plates. A= atrichoblast, T=trichoblast. **(B)** Brightfield images of Col-0 wild type, *kin-12ab* and *kin-12a/b, pGL2>>kin-12f* 5 day old seedlings. **(C)** CLSM section of primary root meristematic epidermal cells in the *kin12a/b, pGL2>>kin-12f* background counterstained with FM4-64. White arrows indicate incomplete crosswalls, yellow arrow indicates amorphous structures of unconfirmed identity. White arrowhead indicates incomplete crosswall highlighted in (D). **(D)** CLSM orthogonal projection of incomplete crosswall indicated by arrowhead in (C). **(E)** CLSM section of primary root meristematic epidermal cells in in the *kin-12a/b x* YFP:RAB-A2a*, pGL2>>kin-12f* background counterstained with FM4-64. White arrows indicate incomplete crosswalls, yellow arrow indicates amorphous structures of unconfirmed identity. **(F-H**) CLSM sections of primary root meristematic epidermal cells expressing YFP:RAB-A2a in Col-0 wild type (F), *kin-12a/b* (G) and *kin-12a/b pGL2>>12f* (H) backgrounds, counterstained with FM4-64. Arrowhead indicates RAB-A2a-positive compartment. **(I,J)** Mean fluorescence intensity of YFP:RAB-A2a orthogonal to a cell plate in a Col-0 wild type, *kin-12a/b,* and *kin-12a/b pGL2>>kin-12f* background. N=41 (*kin-12a/b pGL2>>kin-12f*), 44 (Col-0) and 52 (*kin-12a/b pGL2>>kin-12f*). There is a significant difference in YFP:RAB-A2a distribution in *kin-12a/b* compared to Col-0, and in *kin-12a/b pGL2>>kin-12f* compared to Col-0 and *kin-12a/b* (two-way ANOVA and post-hoc Tukey test, n.s. = p≥0.05; ***=p<0.001). Scale bars, 10µm.

YFP:Kin-12F fluorescence in dividing cells in atrichoblast cell files was abolished, while YFP:Kin-12F was detected at the midzone of dividing cells in trichoblast cell files where *pGL2::kin-12f:cas9-nls-mturquoise* was not expressed (Figure 2A). We concluded that *pGL2::kin-12f::cas9-nls-mturquoise* could target *Kin-12F* and induce KO mutations specifically in atrichobast cell files. We next expressed *pGL2::kin-12f::cas9-nls-mturquoise* in the *kin12a/b* background, and found that 14 of 19 independent transgenic T2 lines produced small, stunted plants (Figure 2B, Figure S3A). Primary roots of these lines showed extensive cytokinesis defects, ranging from misaligned division planes, to incomplete cross-walls, to multi-nucleate cells with no sign of cross-wall formation (Figure 2C,D, S3B,C). We conclude that the three Class II Kin-12s of *Arabidopsis* are essential for cytokinesis in somatic cells and function redundantly.

### Class II Kin-12s are involved in vesicle targeting to the midzone via their interaction with RAB-A2a

When we expressed *pGL2::kin-12f::cas9-nls-mturquoise*, we often observed large amorphous structures that were labelled by FM4-64 (Figure 2C,D, S3B,C) and typically positioned close to the end of incomplete cross-walls. We hypothesised that these may represent mis-targeted membrane material of TGN/EE origin. To investigate further the role of Class II Kin-12s with respect to membrane targeting, we expressed the *pGL2::kin-12f::cas9-nls-mturquoise* construct in a *kin12a/b YFP:RAB-A2a* background. We found that these large amorphous structures were also labelled by YFP:RAB-A2a (Figure 2E), corroborating that they may represent mis-targeted TGN/EE membrane material. We also investigated the impact of loss of Class II Kin-12 function on RAB-A2a distribution at the midzone. In *kin-12a/b YFP:RAB-A2a* plants, YFP:RAB-A2a was still midzone-localised, although quantitative analyses of YFP:RAB-A2a in *kin-12a/b* vs wild type background revealed a small but significant reduction in relative enrichment at the midzone (Figure 2F,G,I; 5.06±0.83 vs 5.37±0.87, respectively). In *kin-12a/b pGL2::kin-12f::cas9-nls-mturquoise* plants, this effect was significantly enhanced (Figure 2F,G,H; relative YFP:RAB-A2a enrichment at the midzone 4.01±0.87), and we observed additional YFP:RAB-A2a punctae in the vicinity of the cell plate (Figure 2F,G,H).

These observations are compatible with a previously unrecognised role of *Arabidopsis* Class II Kin-12s in vesicle transport towards the midzone, possibly mediated by their interaction with RAB-A2a. To test this hypothesis, we expressed and purified the motor domains of Kin-12A, -B and -F from *E.coli* cultures (Figure 3D) and performed *in vitro* gliding assays to test motor activity^39^. In our experiments, both Kin-12A and Kin-12B, but not Kin-12F demonstrated motor activity (Figure 3A-C, E-G). We quantified motor velocities from gliding assays and found Kin-12B velocities to be significantly higher than Kin-12A velocities (0.080±0.011µm/s vs 0.031±0.008µm/s, respectively; Figure 3H). Although motor activity has not been reported for Kin-12A/-12B in *Arabidopsis* before, these results are consistent with recent observations of homologous proteins in *Physcomitrium patens,* where Class II Kin-12s are believed to transport TGN-derived vesicles towards the midzone^21^. In *Arabidopsis*, stable targeting of Kin-12A to the midzone has previously been shown to depend on the midzone-localised microtubule crosslinker MAP65-3^5^. We therefore tested whether MAP65-3 is also required for midzone localisation of Kin-12F by expressing YFP:Kin-12F in the *dyc283/map65-3* mutant, which carries a T-DNA insertion in the *MAP65-3* locus that disrupts protein function^40^. Interestingly, in contrast to what has previously been shown for Kin-12A, we found that YFP:Kin-12F continued to localise to the phragmoplast midzone in the *dyc283/map65-3* background (Figure 3I-L). Taken together, our data demonstrate that Class II Kin-12s are essential for somatic cytokinesis, and contribute towards targeting of YFP:RAB-A2a-positive membranes to the phragmoplast midzone. However, our data also show some *in vitro* and *in planta* differences between Class II Kin-12 members with respect to motor activity and upstream regulators, suggesting their role in cytokinesis may be collaborative rather than identical.

**Figure 3:**
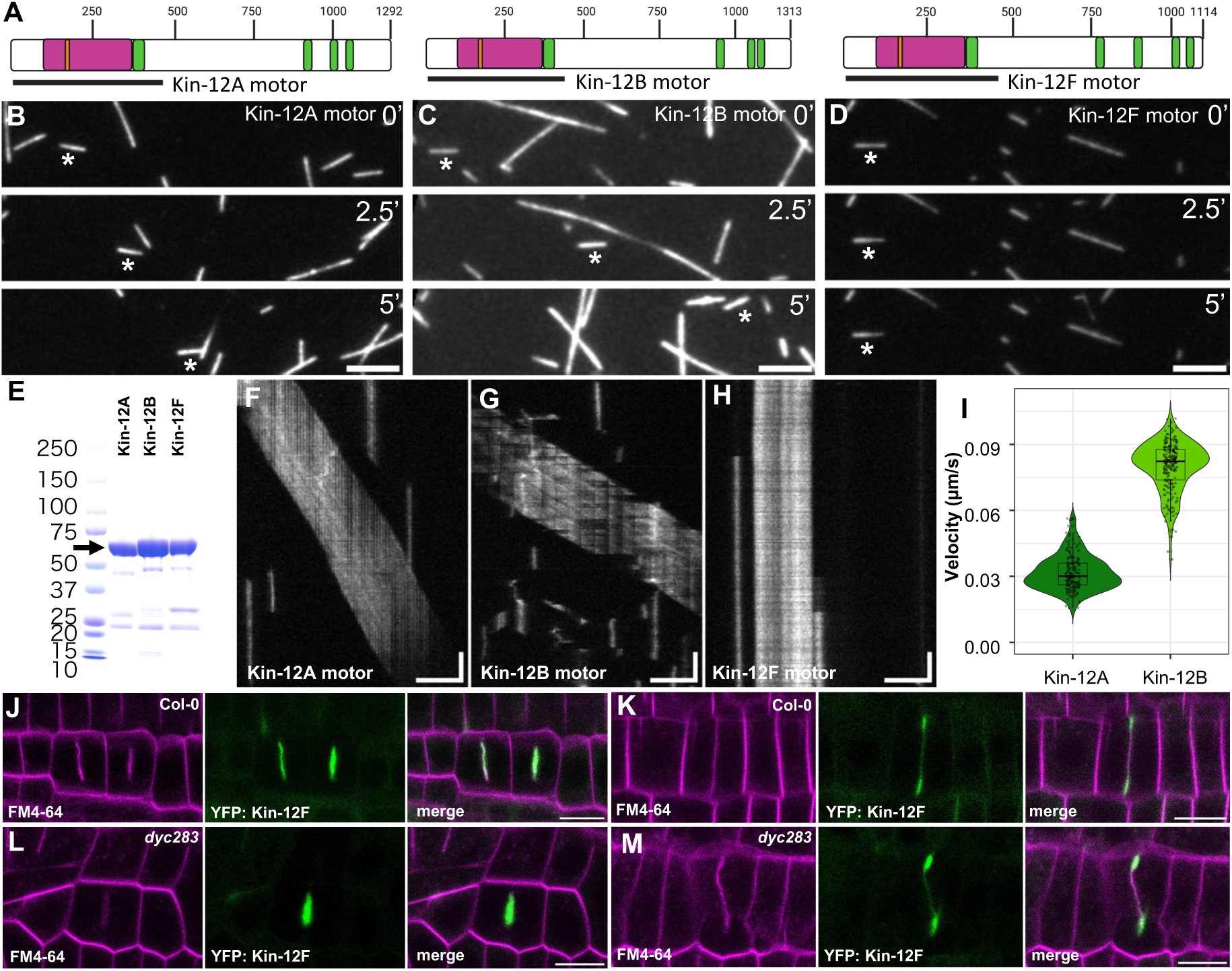
Functional differences between Class II Kin-12 proteins *in vitro* and *in planta*. **(A)** Schematic depiction of Class II Kin-12 proteins with motor domain truncations highlighted by black lines. **(B-D)** TIRF images of Alexa Fluor-568 –labelled GMPCPP-stabilised microtubules with Kin-12A motor domain (B), Kin-12B motor domain (C) and Kin-12F motor domain (D). Asterisks indicate same microtubule tracked over time. **(E)** Coomassie-stained gel of Kin-12A, Kin-12B, and Kin-12F motor domains used in gliding assays as shown in (B-D), (F-H). Arrow indicates bands corresponding to kinesin motor domains. **(F-H)** Kymographs of microtubule gliding assays in the presence of the Kin-12A, -12B, or 12F motor domain (B-D). Note MT gliding activity was observed for Kin-12A and -12B, but not -12F. **(I)** Violin plots of Kin-12A and -12B motor domain velocities calculated from gliding assays shown in (A-C). N = 180 (Kin-12A) and 224 (Kin-12B). **(J-M)** CLSM section of primary root meristematic epidermal or cortical cells in Col-0 wild type (J,K) and *dyc283* (*map65-3*, L,M) expressing *YFP:Kin-12F*, counterstained with FM4-64. Scale bars, 10µm. Graphics created with biorender.com.

### Class II Kin-12s are required for phragmoplast expansion

Following previous reports of a structural role in phragmoplast organisation of Class II Kin-12s in microspores^3^, we also examined phragmoplast morphology and dynamics in somatic cells of Class II Kin-12s mutants. In primary roots of *kin-12a/b* plants expressing RFP:TUB6, phragmoplasts were morphologically indistinguishable from those in wild type plants during disk phase, and were similar to wild type during ring phase apart from a slight reduction in ring-phase phragmoplast width (Figure 4A,B, S4A,B). Surprisingly, we found that phragmoplast expansion rate was over 30% slower in *kin-12a/b* compared to wild-type plants (Figure 4C-E, 0.40±0.07 vs 0.58±0.13µm/min). Phragmoplasts formed normally in *kin-12f RFP:TUB6* plants and were morphologically indistinguishable from those in wild-type plants (Figure 4F,G, S4C,D), but phragmoplast expansion rate was also reduced by 22% compared to wild-type plants (Figure 4H-J, 0.44±0.11 vs 0.57±0.16µm/min). Despite these reductions in phragmoplast expansion rates, primary roots of *kin12a/b* and *kin12f* mutants showed no signs of cytokinesis defects (Figure S3C). We conclude that Class II Kin-12s play a critical role in sustaining phragmoplast expansion, raising the question of whether this effect was coupled to their role in vesicle targeting to the midzone.

**Figure 4:**
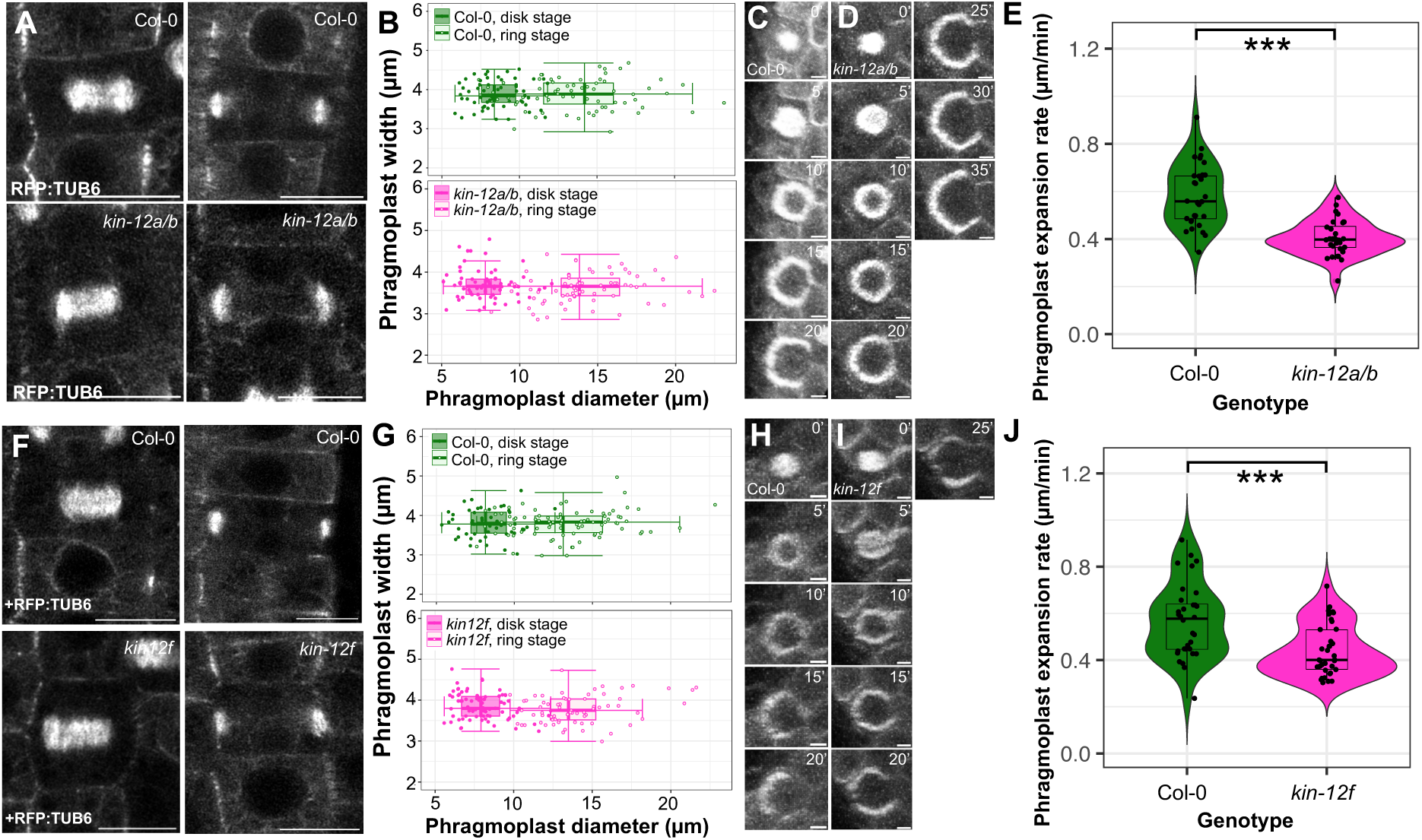
Class II Kin-12 proteins are required for phragmoplast expansion. **(A)** Airyscan CLSM section of primary root meristematic epidermal cells in Col-0 and *kin-12a/b* backgrounds expressing *RFP:TUB6* during disk phase (left) and ring phase (right). **(B)** Box plots of phragmoplast width and diameter in primary root meristematic epidermal and cortex cells in Col-0 and *kin-12a/b* backgrounds expressing *RFP:TUB6* during disk and ring phragmoplast phase. N = 66 (Col-0 disk), 67 (*kin-12a/b* disk), 73 (Col-0 ring), and 79 (*kin-12a/b* ring). **(C,D)** CSLM maximum intensity projections of orthogonally resliced images of primary root meristematic epidermal cells in Col-0 (C) and *kin-12a/b* (D) backgrounds expressing *RFP:TUB6*. Images were taken at 5-minute intervals. **(E)** Violin plots of phragmoplast expansion rates in primary root meristematic epidermal and cortex cells in Col-0 and *kin-12a/b* backgrounds expressing *RFP:TUB6*. Expansion rate was significantly reduced in the *kin-12a/b* background compared to WT (p = 1.202e-08, Student’s T test). *N*=32 phragmoplasts for each genotype. **(F)** Airyscan CLSM section of primary root meristematic epidermal cells in Col-0 and *kin-12f* backgrounds co-expressing *RFP:TUB6* during disk phase (left) and ring phase (right). **(G)** Box plots of phragmoplast width and diameter in primary root meristematic epidermal and cortex cells in Col-0 and *kin-12f* backgrounds expressing *RFP:TUB6* during disk and ring phragmoplast phase. N = 49 (Col-0 disk), 68 (*kin-12f* ring), 75 (*kin-12f* disk), and 83 (Col-0 ring). **(H,I)** CSLM maximum intensity projections of resliced images of primary root meristematic epidermal cells in Col-0 (H) and *kin-12f* (I) backgrounds expressing *RFP:TUB6*. Images were taken at 5 minute intervals. (J) Violin plots of phragmoplast expansion rates in primary root meristematic epidermal and cortex cells in Col-0 and *kin-12f* backgrounds expressing *RFP:TUB6*. Expansion rate was significantly reduced in the *kin-12f* background compared to WT (p = 0.0004159, Student’s T test). *N*=32 phragmoplasts for Col-0, 37 for *kin-12f*.

### RAB-A2a-dependent membrane trafficking is required for proper phragmoplast expansion

To understand whether the role of Class II Kin-12s in phragmoplast expansion was linked to their interaction with functional RAB-A2a, we examined phragmoplast morphology and dynamics when RAB-A2a activity or patterning was perturbed. We first quantified the effect of an inducibly-expressed dominant-negative RAB-A2a[NI] variant (*AtRPS5a>>Dex>>RAB-A2a[NI]*). The morphology of both disk and ring phase phragmoplasts, as quantified by phragmoplast width and diameter, was not affected in our experimental conditions (Figure 5A,B, S4E,F). However, we found phragmoplast expansion rates were reduced by 56% in the presence of RAB-A2a[NI] compared to controls (0.23±0.10µm/min vs 0.53±0.11µm/min; Figure 5C,D,E). We also tested the effect of YFP:Kin-12F-tail expression on phragmoplast expansion, as YFP:Kin-12F-tail expression altered mCh:RAB-A2a distribution during both interphase and cytokinesis (Figure 1, S2). YFP:Kin-12F-tail expression reduced phragmoplast expansion by 28% compared to controls (0.40±0.11µm/min vs 0.55±0.13µm/min; Figure 5H-J). Phragmoplast morphology was largely normal upon by *YFP:Kin-12F-tail* expression, with the exception of ring-phase phragmoplast width, which was slightly increased compared to in control plants (Figure 5F,G, S4G,H). These data indicate that active RAB-A2a at the midzone is required for normal rates of phragmoplast expansion during cytokinesis, linking vesicle delivery to phragmoplast expansion.

**Figure 5:**
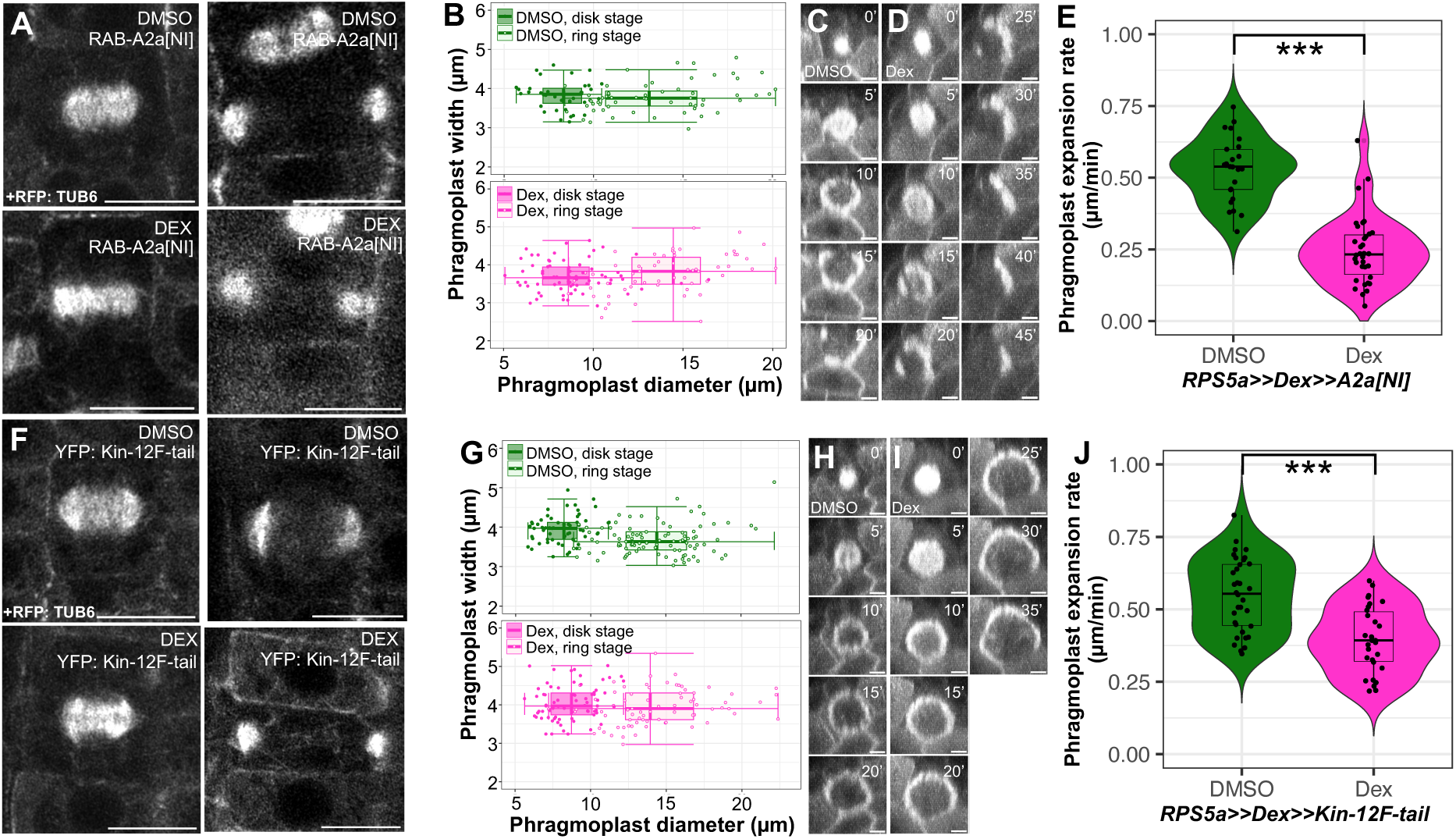
RAB-A2a regulates phragmoplast expansion rates. **(A)** Airyscan CLSM section of primary root meristematic epidermal cells co-expressing *RFP:TUB6* and *DEX>> RAB-A2a[NI]* upon 36hr treatment with 500nM Dex or DMSO during disk phragmoplast phase (left) and ring phragmoplast phase (right). **(B)** Box plots of phragmoplast width and diameter in primary root meristematic epidermal cells co-expressing RFP:TUB6 and DEX>>RAB-A2a[NI] upon 36hr treatment with DMSO (top) or 500nM Dex (bottom) during disk and ring phragmoplast phase. N = 31 (DMSO disk), 43 (DMSO ring), 50 (Dex ring), and 57 (Dex disk). **(C,D)** CSLM maximum intensity projections of resliced images of primary root meristematic epidermal cells co-expressing RFP:TUB6 and DEX>>RAB-A2a[NI] upon 36hr treatment with DMSO (C) or 500nM Dex (D). Images were taken at 5 minute intervals. **(E)** Violin plots of phragmoplast expansion rates in primary root meristematic epidermal and cortex cells co-expressing RFP:TUB6 and DEX>>RAB-A2a[NI] upon 36hr treatment with DMSO or 500nM Dex. Expansion rate was significantly reduced in Dex-compared to DMSO-treated plants (p = 6.93e-13, Student’s T test). *N*=25 (DMSO) and 34 (Dex). **(F)** Airyscan CLSM section of primary root meristematic epidermal cells co-expressing RFP:TUB6 and DEX>>YFP: Kin-12F-tail upon 72hr treatment with 5µM Dex or DMSO during disk phragmoplast phase (left) and ring phragmoplast phase (right). **(G)** Box plots of phragmoplast width and diameter in primary root meristematic epidermal cells co-expressing RFP:TUB6 and DEX>>YFP:Kin-12F-tail upon 72hr treatment with DMSO (top) or 5µM Dex (bottom) during disk and ring phragmoplast phase. N = 57 (DMSO disk), 73 (Dex disk), 78 (Dex ring), and 97 (DMSO ring). **(H,I)** Resliced CSLM maximum intensity projections of primary root meristematic epidermal cells co-expressing RFP:TUB6 and DEX>>YFP:Kin-12F-tail upon 72hr treatment with DMSO (H) or 5µM Dex (I). Images were taken at 5 minute intervals. **(J)** Violin plots of phragmoplast expansion rates in primary root meristematic epidermal and cortex cells co-expressing RFP:TUB6 and DEX>>YFP:Kin-12F-tail upon 72hr treatment with DMSO or 5µM Dex. Expansion rate was significantly reduced in in Dex-compared to DMSO-treated plants (p = 3.469e-06, Student’s T test). *N*=34 (DMSO), 30 (Dex). Scale bars, 10µm (A&F) or 5µm (C,D,H,I).

### The Kin-12A/-12B interactor TIO kinase is required for phragmoplast formation in somatic cells

Kin-12A and -12B have previously been shown to interact with TWO-IN-ONE (TIO), which is required for proper cytokinesis in both reproductive and somatic cells of *Arabidopsis*^16^. TIO is the sole *Arabidopsis* orthologue of the sonic hedgehog pathway component known as Fused kinase in *Drosophila* and Stk36 in humans, which are involved in control of cell proliferation and differentiation^26^. While the exact function and substrates of TIO kinase in plants remain unknown, it has been identified as a positive regulator of phragmoplast expansion in *Arabidopsis* microspores^17^. We therefore investigated whether the roles of Class II Kinesin-12 proteins and RAB-A2a in phragmoplast expansion during somatic cell cytokinesis were linked to TIO kinase function. We first investigated in greater detail the role of TIO kinase during somatic cell cytokinesis. Homozygous knockout mutants of TIO kinase have previously been shown to be unobtainable due to germline sterility^16,17^. We therefore used CRISPR-Cas9 driven from the GL2 promoter to target TIO kinase in somatic cells (*pGL2::tio::cas9-nls-mturquoise)*. To confirm whether the *pGL2::tio::cas9-nls-mturquoise* construct was able to efficiently knock out TIO kinase, we first introduced it into lines inducibly expressing YFP:TIO (*AtRPS5a>>DEX>>YFP:TIO*). In the absence of *pGL2::tio::cas9-nls-mturquoise,* YFP:TIO localised to the phragmoplast midzone, and followed the expanding phragmoplast as cytokinesis progressed (Figure S5A&B). This pattern is consistent with previous immunolocalisation of TIO kinase in cultured cells of *Arabidopsis*^16^. However, this signal was abolished in atrichoblast cell files of *pGL2::tio::cas9-nls-mturquoise,* indicating that our construct could knock out TIO in a tissue specific manner as intended (Figure S5C). In the T1 generation of all backgrounds in which we expressed *pGL2::tio:cas9-nls-mturquoise* plants, we recovered plants that were extremely stunted (Figure 6A). These plants had numerous cytokinesis defects in their primary roots, including the presence of incomplete cross-walls in some cells, as well as large, multinucleate cells that showed no signs of having undergone cytokinesis (Figure 6B). As we had previously observed in *pGL2::kin-12f:cas9-nls-mturquoise* plants, cytokinetic failure was sometimes, albeit less frequently, accompanied by the appearance of amorphous structures (Figure 6B).

**Figure 6:**
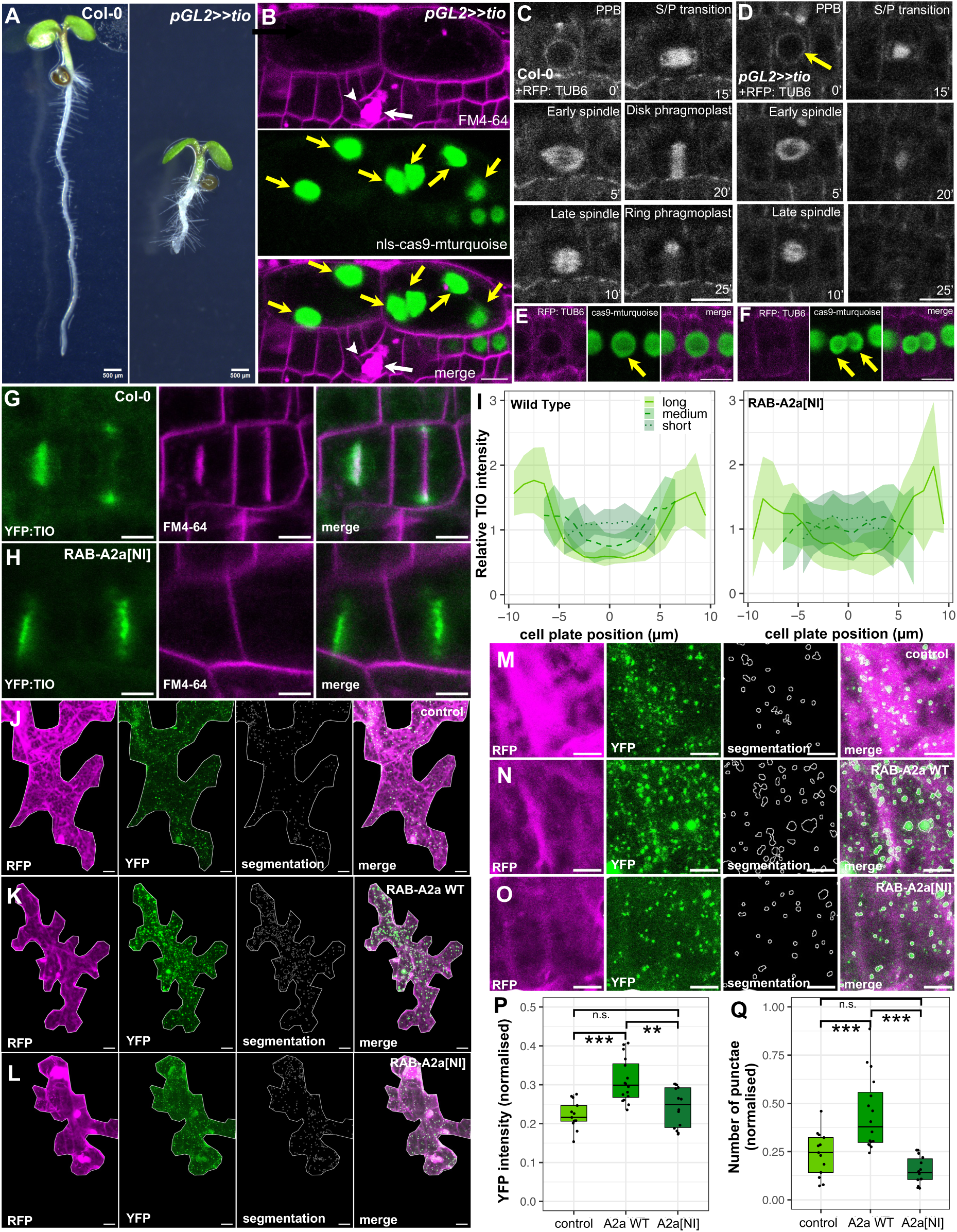
TIO localization at the midzone is enhanced by RAB-A2a-Kin-12A interactions. **(A)** Brightfield images of Col-0 and *pGL2>>tio pGL2::tio::cas9-nls-mturquoise* 5 day old T1 seedlings. **(B)** CLSM section of primary root meristematic epidermal cells in the *pGL2::tio::cas9-nls-mturquoise (pGL2>>tio)* background counterstained with FM4-64. White arrowhead indicates incomplete crosswalls, white arrow indicates amorphous structures of unknown identity, yellow arrows indicate nuclei (cas9 expression) in multinucleate cells. **(C,D)** CLSM sections of primary root meristematic epidermal cells of Col-0 background (C) or *pGL2>>tio* background (D) expressing *RFP: TUB6*. Imaged at 5 min intervals. Yellow arrow indicates nucleus monitored in (E) and (F). **(E,F)** CLSM sections of primary root meristematic epidermal cells of *pGL2>>tio* background before (E) and after (F) mitosis. **(G,H)** CLSM sections of primary root meristematic epidermal cells expressing Dex>>YFP: TIO in a wild type or Dex>>RAB-A2a[NI] background. Roots were stained with FM4-64. **(I)** Relative fluorescence intensity of YFP:TIO along cell plates as those shown in (G,H). Cell plates were grouped by diameter into short (<9µm), medium (9-13µm) and long (>13µm). Lines are mean values, shaded areas are +/- 1SD. N = 10 (Dex>>RAB-A2a[NI] long), 16 (Col-0 medium, Col-0 long), 17 (Dex>>RAB-A2aNI medium), 18 (Col-0 short), and 34 (Dex>>RAB-A2a[NI] short). **(J-L)** Segmented *Nicotiana benthamiana* epidermal leaf cells transiently expressing *nYFP:TIO C-terminus* with *cYFP: Kin-12A tail* and *mRFP1* from a ratiometric rBiFC vector alone at O.D. 600 0.1 (J) or alongside wild-type RAB-A2aWT (K), or RAB-A2aNI (L) at O.D. 600 0.1 (combined O.D. 600 of 0.2). **(M-O)** cropped sections from cells as those shown in (J-L). **(P,Q)** Quantification of mean YFP intensity (P) or number of punctae per area (Q) normalised against mean RFP fluorescence from images as those shown in (J-L). Note the presence of RAB-A2aWT, but not RAB-A2aNI significantly increased both YFP intensity and number of punctae. Cell area, mean RFP intensity, or mean intensity of punctae did not significantly differ between conditions (one-way ANOVA and post-hoc Tukey test, n.s. = p≥0.05; **=p<0.01 ;***=p<0.001). N= 13 (control), 14 (A2aNI), and 16 (A2aWT). Scale bars, 500µm (A), 10µm (B,J-K), 5µm (C-H,M-O).

We next used *pGL2::tio::cas9-nls-mturquoise* in the RFP:TUB6 background to monitor the effect of TIO suppression on phragmoplast dynamics. Surprisingly, we observed that of 25 nuclear divisions imaged in RFP:TUB6 *pGL2::tio::cas9-nls-mturquoise* primary root epidermal cells, 13 ended with complete failure to form a recognisable phragmoplast following spindle dissolution (Figure 6C,D). In place of the phragmoplast, these cells often contained amorphous tubulin aggregates at the cell centre (Figure S5D) and failed to form cross-walls, resulting in bi- or multi-nucleate cells (Figure 6E,F). We also noted that YFP: RAB-A2a signal failed to accumulate at the midzone in *pGL2::tio::cas9-nls-mturquoise* lines (Figure S5E) indicating that TIO plays a critical role in phragmoplast formation as well as expansion.

### Class II Kinesin-12 and RAB-A2a are required for proper TIO kinase patterning

We next tested whether Class II Kin-12s and RAB-A2a are required for proper TIO kinase localisation during cytokinesis. We first examined whether like Kin-12A and -12B, Kin-12F also interacts with TIO. Comparative alignments of the amino acid sequences of the Kin-12A,-12B, and -12F tail regions show that much of the TIO interacting domain conserved in the Kin-12A and -12B tail regions^16^ is absent in Kin-12F (Figure S6A). In line with this, we could find no evidence for an interaction between the Kin-12F tail region and TIO in Y2H (Figure S6B,C), although we were able to reproduce the previously reported interaction between full-length TIO and its C-terminus with the tail regions of Kin-12A and -12B^16^ (Figure S6B,C). We therefore focussed our analyses of YFP:TIO localisation on the *kin12a/b* mutants. When expressed in the *kin12a/b* background, YFP:TIO still localised to the phragmoplast midzone and was enriched at the leading edge as in wild type plants (Fig S6D-F). However, we recorded a significant reduction in YFP:TIO at the midzone, but enhanced enrichment in a zone up to 2µm from the midzone, which corresponds to the area occupied by phragmoplast (Fig S6D,E,N). This is consistent with a scenario in which Kin-12A and -12B contribute to localising TIO kinase to the phragmoplast midzone alongside other TIO kinase interactors ^18^. To test whether RAB-A2a activity is required for TIO kinase localisation during cytokinesis, we co-expressed YFP:TIO with RAB-A2a[NI]. In the presence of RAB-A2a[NI], YFP:TIO still localised to the phragmoplast midzone and, in contrast to the *kin-12a/b* background, did not become more laterally diffuse (Fig S6O,P). However, YFP:TIO patterning did become significantly more variable along the length of the cell plate in the presence of RAB-A2a[NI] compared to controls (Fig 6G-I, S6L). Such an increase in intensity variation did not occur for midzone-localised GFP:MAP65-3 in the presence of RAB-A2a[NI] (Fig S6G-J,M), indicating that this change in YFP:TIO pattern does not reflect a generic midzone geometry or structure change caused by RAB-A2a[NI] expression. Taken together, these data indicate that both Kin-12A/-12B and RAB-A2a contribute to TIO patterning, albeit in different ways.

### RAB-A2a binding to Kin-12A promotes its interaction with TIO Kinase

To build a functional model for Class II Kin-12s – RAB-A2a – TIO function at the midzone, we fine-mapped the interaction domains between these proteins in Y2H. The TIO interaction domain of Kin-12A and -12B^16^ was included in the Class II Kinesin-12 clones we identified as interacting with RAB-A2a in Y2H (Figure 1A). To increase our resolution of the relative positions of the interaction domains of TIO and RAB-A2a in the tail regions of Kin-12A and -12B, we expressed truncated variants of these tail regions in Y2H that encompassed either the Kin-12A/-B TIO-Binding Domain (TBD), or the remainder of the original clone without the TIO binding domain (Kin-12A/B-tailΔTBD; Figure S7A,B). In Y2H, neither Kin-12A-TBD nor Kin-12B-TBD interacted with RAB-A2a, its mutant variants, or other Rab GTPases known to localise to the cell plate (Figure S7C,D). However, both Kin-12A-tailΔTBD and Kin-12B-tailΔTBD interacted with RAB-A2a WT and RAB-A2a[QL] in Y2H (Figure S7C,D), as had been observed for the full-length Kin-12A/-12B tail regions (Figure 1A, S7C,D). This demonstrates that the TBD and RAB-A2a binding domains of Kin-12A and -12B are adjacent and non-overlapping. Interestingly, in an equivalent truncated clone of Kin-12F (Kin-12F-tailΔC-term), the binding affinity for RAB-A2a, its mutant variants and other Rab GTPases was increased compared to the full-length Kin-12F tail region (Figure S7C,D).

Based on these observations, we hypothesised that the binding of RAB-A2a to Kin-12A/-12B may affect their interaction with TIO at the adjacent binding site. To test this hypothesis, we used a ratiometric bimolecular fluorescence complementation (rBiFC) vector system^41^. This system allows ratiometric expression of two proteins of interest fused to the N- or C-terminus of a split eYFP, alongside mRFP1 as an expression control all from the same vector^41^. We used this system to transiently express nYFP:TIO-C-terminus with either cYFP:Kin-12A-tail or cYFP:Kin-12F-tail in *Nicotiana benthamiana* leaves. We consistently observed YFP punctae in cells co-expressing nYFP:TIO-C-terminus and cYFP:Kin-12A-tail (Fig 6J,M), but never in cells co-expressing nYFP:TIO C-terminus and cYFP:Kin-12F-tail (Fig S8D-I). We manually segmented cells from maximum intensity projections and quantified mean YFP fluorescence, YFP punctae per area, and RFP fluorescence in cells co-expressing nYFP:TIO-C-terminus and cYFP:Kin-12A-tail (Figure S8A-C). As there was a correlation between both mean YFP fluorescence and RFP fluorescence (R² = 0.5259), as well as YFP punctae per area and RFP fluorescence (R² = 0.4062), we normalised mean YFP fluorescence and YFP punctae per area by mean RFP fluorescence per cell to account for diberences in rBiFC expression levels (Fig 6P,Q).

To test whether expression of *RAB-A2a* had an ebect on the TIO – Kin-12A interaction, we co-infiltrated wild-type RAB-A2a and RAB-A2a[NI] variants with the rBiFC construct. We found that co-expression of wild-type RAB-A2a significantly increased both mean YFP fluorescence and numbers of YFP punctae per cell (Fig 6K,N,P,Q). This was not the case when we co-expressed RAB-A2a[NI] (Fig 6L,O,P,Q), indicating that (1) the presence of RAB-A2a could enhance the interaction between TIO and Kin-12A *in vivo* and (2) this enhancement was dependent on RAB-A2a activity. These observations are consistent with a scenario in which RAB-A2a binding aids TIO recruitment to Kin-12A/-12B at the midzone to initiate phragmoplast remodelling, thus coupling vesicle delivery and phragmoplast remodelling during cytokinesis.

## Discussion

In this study, we demonstrate that the interaction between the membrane-associated Rab-A GTPase RAB-A2a and phragmoplast-associated Class II Kin-12s plays an essential role during cytokinesis in *Arabidopsis*. Successful cytokinesis in plants depends on the addition of vesicles from the TGN/EE to the cell plate, and spatio-temporal co-ordination of phragmoplast dynamics with this. Our data suggests that the interaction between RAB-A2a and Class II Kin-12 is essential for both of these processes: first, we have shown that Class II Kin-12s are required for somatic cytokinesis, during which they position RAB-A2a-associated membranes correctly at the phragmoplast midzone, and secondly, we have demonstrated that both Class II Kin-12s and functional RAB-A2a are required for phragmoplast expansion at physiological rates. Through co-operatively platforming TIO kinase at the midzone, we propose that this Class II Kin-12-RAB-A2a module acts to couple phragmoplast polymerisation to vesicle trafficking rates.

The question of exactly how Class II Kin-12s contribute to targeting of membranes at the phragmoplast midzone remains open: both Kin-12A and -12B were motile in *in vitro* gliding assays, indicating that they may play a role in vesicle transport to the midzone. However, we did not observe motility for Kin-12F. Whilst we cannot definitively conclude based on this that Kin-12F does not have motility *in planta,* it is noteworthy that Kin-12F is the only Kin-12 protein that does not have a conserved proline-tyrosine (PY) motif in its motor domain, which have been implicated in control of motor activity in other kinesins^37^. The absence of Kin-12F from the phragmoplast leading edge, the main site of membrane delivery, also argues against a role in vesicle transport. We recorded slightly depleted RAB-A2a midzone targeting in *kin-12a/b* backgrounds compared to wild-type backgrounds, an effect that was enhanced when all three Class II Kin-12s were lost. We also often observed large, amorphous structures that were labelled by both RAB-A2a and FM4-64 when all three Class II Kin-12s were lost, which were frequently positioned close to the end of incomplete cross-walls. While it is therefore clear that Class II Kin-12s are required for proper midzone targeting of RAB-A2a-associated membranes during cytokinesis, it is difficult to discriminate whether this loss of targeting in the absence of Class II Kin-12s arises from failure to platform RAB-A2a-bearing vesicles at the midzone after their delivery, and/or a failure to transport RAB-A2a-bearing vesicles to the midzone along the phragmoplast. Both or a combination of these scenarios could lead to a situation in which mis-targeted vesicles fuse to form membrane structures away from the midzone, thus preventing proper cell plate formation.

One possibility is that Kin-12A/B and -F fulfil transport and platforming functions to different degrees: we found that localisation of Kin-12F to the midzone is not dependent on the interdigitating microtubule crosslinker MAP65-3, as has been shown previously for Kin-12A^5^. Kin-12F is depleted at the leading edge of the cell plate/phragmoplast, but is present at the lagging transition zone where the cell plate becomes a tubulo-vesicular network. Here, the majority of microtubules attach to the cell plate without the participation of MAP65-3. We therefore propose that Kin-12F may act at the transition zone to promote targeting of vesicles during the formation of the cell plate tubulo-vesicular network, after their initial delivery to the phragmoplast leading edge. As we could find no evidence that Kin-12F interacts with TIO kinase, it is also possible that Kin-12F acts from this position as a “buffer” that can accumulate RAB-A2a membranes without triggering TIO kinase recruitment and further phragmoplast expansion. This implies the relative abundance of Kin-12A,B, and F may fine-tune cytokinesis dynamics in different cell types, which could account for the phenotypic differences observed between somatic and germline divisions^17,18^.

RAB-A2a interacts with Kin-12A and -12B in an adjacent, but non-overlapping domain, to that of TIO kinase. We have also shown that Class II Kin-12s and RAB-A2a are required for proper TIO kinase patterning during cytokinesis *in planta,* and that presence of functional RAB-A2a can enhance the interaction of TIO kinase and Kin-12A *in vivo.* TIO kinase has previously been shown to be a positive regulator of phragmoplast expansion in microspores, but our data suggest that it is also more broadly required for phragmoplast formation^16,17^. We found that in somatic cells, suppression of TIO kinase causes failure to form phragmoplasts after spindle dissolution in approximately 50% of divisions. This suggests that TIO kinase is required for phragmoplast formation, as well as expansion, in somatic cells. TIO kinase is the sole *Arabidopsis* homologue of Fused kinase/Stk36, which is a component of the sonic hedgehog pathway in metazoans^16^. No other components of the sonic hedgehog pathway appear to be conserved in land plants, and while the targets of TIO kinase remain unidentified, they have been suggested to include Class II Kin-12s themselves^17^. It has recently been shown that the chemical compound PP2 blocks phragmoplast formation in a manner strikingly similar to *RFP:TUB6 pGL2::tio:cas9-nls-mturquoise* mutants^42^. The authors identified that PP2 treatment disrupted phosphorylation of Kin-12A, and proposed that PP2 may block phragmoplast formation by inhibiting phosphorylation of Class II Kin-12s^42^. Combining these observations with those presented here, it is tempting to speculate that PP2 treatment may block TIO kinase activity. This could explain the similar effects on phragmoplasts caused by PP2 treatment and *pGL2:: tio: cas9-nls-mturquoise* expression. It is not entirely clear how TIO kinase-driven phosphorylation of Class II Kin-12s could contribute to phragmoplast expansion . Recent data suggest that a Kin-12A variant with reduced phosphorylation is excessively stabilised on microtubules^42^, which could be inhibitory to microtubule turnover required for phragmoplast growth. However, we did not observe any defects in transition from solid to disk-shaped phragmoplasts, or other substantial defects in phragmoplast morphology, when either Kin-12s or RAB-A2a function was impaired, suggesting that microtubules were not excessively stabilised. To account for these observations, we instead propose that TIO kinase acts to promote polymerisation and stabilisation of microtubules of the phragmoplast. This can account for our observation that TIO kinase is involved in both phragmoplast formation after spindle dissolution as well as its later expansion. It is not clear whether this function emerges directly from Kin-12A/B phosphorylation, but TIO kinase may also have other, yet unknown targets that contribute to phragmoplast remodelling.

Based on our data, we propose the following model for the function of a RAB-A2a/Kin-12/TIO module in spatio-temporal control of cytokinesis: (1) RAB-A2a-bearing vesicles are brought to the midzone along phragmoplast microtubules by transport kinesins, which include Kin-12A/B. (2) Kin-12A/B associate with microtubule-crosslinker MAP65-3 to stably platform vesicles at the leading edge, thus spatially patterning initial vesicle tethering and fusion. (3) Kin-12F is absent from the cell plate leading edge, but is enriched at the transition zone, where it may act as a scaffold to stabilise microtubule-membrane interactions independently of MAP65-3 promoting formation of a tubulo-vesicular network and eventually a stable cell plate. (4) The interaction of RAB-A2a with the tail regions of Kin-12A/-12B promotes recruitment of TIO kinase to the adjacent part of the Kin-12A/-12B tail regions, possibly via triggering a conformational change, therefore driving TIO kinase localisation to the midzone proportionally to RAB-A2a abundance at the midzone. (5) TIO activity promotes phragmoplast microtubule polymerisation, allowing first phragmoplast formation, and later phragmoplast expansion (Figure 7). This model provides a testable, mechanistic explanation for previous observations regarding the coupling of vesicle delivery and phragmoplast growth^31–33^, and explains how plants may ensure uniform cell plate morphology during cytokinesis.

**Figure 7:**
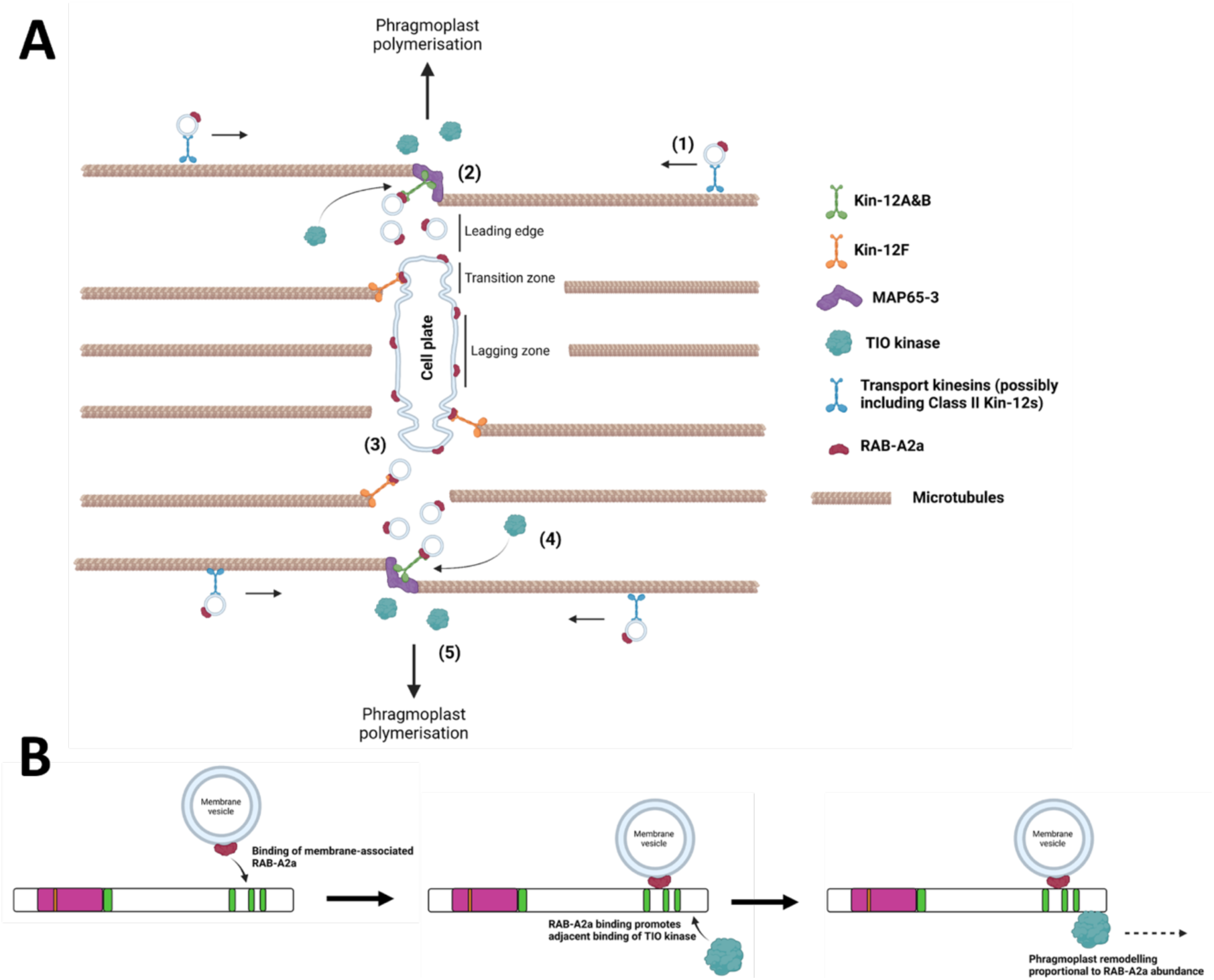
Model for Class II Kin-12-RAB-A2a-TIO module function during cytokinesis. **(A)** (1) Transport of RAB-A2a-bearing vesicles to the midzone is accomplished at least partially by kinesins of uncertain identity, but Kin-12A&B may play a role in this. (2) Kin-12A (and likely Kin-12B) is stabilised at the leading edge phragmoplast midzone by the microtubule crosslinker MAP65-3, where they provide a platform for incoming RAB-A2a-bearing vesicles. (3) Kin-12F provides an additional platform at the transition zone midzone that is independent of MAP65-3, supporting the formation of the tubulo-vesicular cell plate (4) RAB-A2a activity and Kin-12A&B dictate proper TIO kinase localisation at the phragmoplast midzone, which (5) couples stable phragmoplast polymerization to membrane presence at the midzone. **(B)** Model for recruitment of RAB-A2a and TIO kinase to the midzone. Binding of membrane-associated RAB-A2a to the Kin-12A&B tail regions promotes binding of TIO kinase to the adjacent part of the tail regions, possibly through inducing a steric change in the kinesin tail region. TIO kinase activity at the midzone then promotes phragmoplast remodelling proportional to RAB-A2a abundance at the midzone. Graphic created with biorender.com.

## Materials and Methods

### Plant materials, growth and treatments

The *Arabidopsis thaliana* ecotype Columbia was used throughout with the exception of the *dyc283* mutant of MAP65-3, which is in Wassilewskija ecotype^40^. *YFP: Kin-12F dyc283* plants described in Figure 3 are therefore Columbia - Wassilewskija hybrids. Kin-12A is denoted as At4g14150 in the *Arabidopsis* genome, Kin-12B as At3g23670, and Kin-12F as At3g20150. The transgenic lines *pRAB-A2a::YFP:RAB-A2a*^25^, *35S>>DEX>>RAB-A2a[SN]*^25^, *p35s::mCherry:RAB-A2*a^30^, *kinesin12a/b*^3^, *pUBQ10::RFP:TUB6*^43^ and *proMAP65-3::GFP:MAP65-3*^40^ have all been described before. The *kinesin12f* SALK insertion line described here is SALK_039654C (NASC ID N655459). All plants were grown at 20°C in a 16h:8h day:night cycle. Primary roots were imaged 3-5 days after germination on half-strength Murashige and Skoog medium (MS, Sigma Aldrich) plates with 1% w/v sucrose and 0.8% Difco-agar (Appleton Woods) at pH 5.7. For constructs whose expression was induced conditionally via dexamethasone (Dex), plants were grown for 2.5 days from germination on half-strength solid MS medium before transfer to liquid half-strength MS medium (1% w/v sucrose, pH5.7) for the indicated time period containing either Dex (Sigma Aldrich – diluted from a 20mM stock in DMSO) at the indicated concentration, or the equivalent concentration of DMSO. Plants in liquid half-strength MS were gently agitated on an orbital shaker at 50rpm throughout the induction period. Treatment with Brefeldin-A (Sigma-Aldrich) was performed in water for the indicated time period with 50µM Brefeldin-A (diluted from a 50mM stock in DMSO) or the equivalent concentration of DMSO. For GUS staining, plants were incubated in a solution of 1mM X-Gluc, 0.05% Triton X-100, 5mM EDTA, 50mM sodium phosphate, 0.5mM potassium ferrocyanide and 0.5mM potassium ferricyanide for 1 hour at 37°C. Staining of primary roots was then photographed directly. For staining with FM4-64 (Thermo Fisher Scientific), plants were incubated for 15 minutes in a solution of 0.1µg/mL prior to imaging (diluted from a 1mg/mL stock in water). Transformation of novel transgenes into plants was performed *via Agrobacterium*-mediated floral dip^44^.

### Molecular Cloning

All genes were amplified by PCR with Phusion^TM^ High-Fidelity DNA Polymerase (Thermo Fisher Scientific) from cDNA of *Arabidopsis* ecotype Columbia. *DEX>>RPS5a>>YFP: Kinesin-12F, DEX>>YFP: Kinesin-12F Tail* and *DEX>>RPS5a>>YFP-TIO* were all generated by cloning of the relevant cDNA region into a YFP-containing pENTRY vector via digestion with the restriction endonuclease AscI (New England Biolabs). The resulting YFP-fusion transgenes were then transferred into *pOpIN2-RPS5a*^35^ using Gateway^TM^ LR Clonase II Enzyme Mix (Thermo Fisher Scientific). Tail regions of Kinesin-12A, Kinesin-12B and Kinesin-12F (and truncations of them) used in pairwise Y2H tests were amplified from cDNA with primers containing EcoRI and XhoI restriction endonuclease digest sites and cloned into the vector pJET1.2/blunt (CloneJET, Thermo Fischer Scientific). In the cases of Kinesin-12B & Kinesin-12F, an internal EcoRI site in each was first removed by PCR. The tail regions were excised from pJET1.2/blunt via digestion with EcoRI and XhoI (New England Biolabs) and ligated into the activation-domain vector pAD-Gal4 (Stratagene) using T4 DNA ligase (Thermo Fisher Scientific). Rab GTPases and mutant variants of them used in pairwise Y2H tests were amplified from cDNA or pre-existing templates with primers containing KpnI and SmaI restriction endonuclease digest sites, and cloned into the binding-domain vector pLexA-C (Clonetech), also *via* the intermediate vector pJET1.2/blunt (CloneJET, Thermo Fischer Scientific). All cloned Rab GTPase variants used for Y2H lack the DNA sequence for the final 6 amino acids so as to prevent geranylgeranylation in yeast. For pairwise Y2H tests between Kinesin-12 tail regions and TIO kinase, pGADT7 and pGBKT7 (Clonetech) containing Kinesin-12A&B tail regions and TIO kinase respectively have been described before^17^. For cloning of Kinesin-12F tail regions into pGADT7, Kinesin-12F tail region truncations were amplified from cDNA with primers containing XmaI and XhoI digest sites and introduced into pGEM®-T Easy (Promega). These were then removed via digestion with XmaI and XhoI (New England Biolabs) and ligated into XhoI/XmaI-digested pGADT7 using T4 DNA ligase (Thermo Fisher Scientific).

For CRISPR-Cas9 targeting of Kinesin-12F and TIO, four target sites for each gene were identified using ChopChop (https://chopchop.cbu.uib.no/). Target sites were incorporated into two complementary olignonucleotides prefaced with the sequences *attg* and *aaac* respectively. Each oligonucleotide pair was then hybridised via denaturing at 98°C for 5 minutes and subsequent incubation at room temperature for 5 minutes. Hybridised oligonucleotides were then incorporated into plasmids pDGE5, pDGE7, pDGE9 and pDGE11 respectively^45^ *via* a Golden Gate reaction with the type-IIS restriction endonuclease BpiI (New England Biolabs). These were then simultaneously transferred into a single vector via a Golden Gate level 1 reaction with BsaI (New England Biolabs) and a modified version of pICH47742^46^. For tissue-specific CRISPR from the *GLABRA2* promoter, the final constructs were assembled via a Golden Gate level 2 reaction between pICH47732-FAST-Red (Position 1^47^), pICH47742-oligonucleotides (Position 2), pICH47751-pGL2::zCas9iNLs-2AmturquoiseN7-rnsCE9t (Position 3), pICH41766 (Position 4^46^) and pAGM4673^46^ using BpiI.

For rBiFC vectors containing split-YFP variants of Kin-12A tail, -12F tail and the TIO C-terminus, these regions were amplified from *Arabidopsis* cDNA and introduced into the Gateway vectors pDONR221-P3P2 (-12A tail, -12F tail) and pDONR221-P1P4 (TIO C-terminus) using BP clonase II enzyme mix (Invitrogen/Thermo Fischer Scientific). These were subsequently simultaneously introduced into pBiFCt-2in1-NN^41^ using LR clonase enzyme mix (Thermo Fischer Scientific). RAB-A2a variants co-expressed in *Nicotiana* with rBiFC constructs were cloned from pre-existing templates into the Gateway vector pDONR207 (Invitrogen/Thermo Fischer Scientific) with BP clonase II enzyme mix, and subsequently into pUB-DEST^48^ with LR clonase enzyme mix. All constructs described in this study were validated via Sanger sequencing (Microsynth) and restriction digests. *Escherichia coli* strains DH5α and DB3.1 were used for molecular cloning. Constructs were introduced into the *Agrobacterium tumefaciens* strain GV3101::pMP90 by electroporation prior to transformation of *Arabidopsis*.

### Yeast two-hybrid

For the initial Y2H screen and subsequent pairwise Y2H tests between Kinesin-12 members and Rab GTPases, the haploid NYM51 and NMY61 (Dualsystems Biotech) strains of *Saccharomyces cerevisiae* were used. For the screen, NYM61 cells were transformed with a Gal4-AD-fused *Arabidopsis* cDNA library via lithium acetate-mediated transformation^48^, and subsequently mated with NMY51-expressing pLexA-C RAB-A2aQL. Subsequent selection conditions for positive interactions were as previously described^49^. For later confirmation via independent pairwise Y2H tests, NYM51 was transformed with pLexA-C containing Rab GTPase variants, NMY51 was transformed with pAD-Gal4 containing Kinesin-12 tail regions (see above) and transformants selected on SD-Leu or SD-Tryp medium. Diploids were obtained *via* mating on YPDA medium and subsequent selection on SD-Leu-Tryp medium. For pairwise Y2H tests between Kinesin-12 members and TIO kinase, the diploid strain AH109 (Clonetech) was used to mimic the original conditions identifying this interaction^30^. AH109 was simultaneously transformed with pGADT7-containing Kinesin-12 tail truncations and pGBKT7-containing TIO kinase truncations via lithium acetate-mediated transformation^48^, and dual transformants selected on SD-Leu-Tryp medium. For all interaction tests displayed, diploids were grown in liquid SD-Leu-Tryp medium overnight at 30°C. Cultures were diluted to O.D_600_ of 1, 0.1 and 0.01 in sterile MilliQ water before plating onto SD-Leu-Tryp and SD-Leu-Tryp-His-Ade plates with a multichannel pipette. Plates were then incubated between 2-4 days at 30°C. All pairwise Y2H tests were performed independently at least three times, and a representative result is shown.

### Co-IPs and immunoblots

For testing co-immunoprecipitation between YFP: Kinesin-12F/YFP: Kinesin-12F-tail and mCh:RAB-A2a, a method adapted from that described in^50^ was used. Briefly, plants were grown in liquid ⅓-strength MS (1% w/v sucrose, pH5.7) for 7 days with gentle agitation of 50rpm on an orbital shaker. At 7 days, 20µm Dex was added for 16 hrs. Root and shoot tissue were then harvested and homogenised in an ice-cold isolation buffer with 2% w/v CHAPS (Sigma-Aldrich). The homogenate was then centrifuged at 14,000rpm for 5 minutes at 4°C. The supernatant was removed and 1mM DTSSP added, then incubated on ice for 15 minutes. DTSSP activity was then quenched by addition of 50mM Tris-HCl pH7.5, and a further 5 minutes incubation on ice. The mixture was then centrifuged at 14,000rpm for 10 minutes at 4°C. 50µl of µMACS anti-GFP magnetic beads (Miltenyi Biotec) were added to the supernatant, and this was then incubated for a further 30 minutes on ice. Bead complexes were then washed and isolated using µMACS columns (Miltenyi Biotec) as previously described (Kalde et al., 2019). Immunoblots against YFP: Kinesin-12F and RAB-A2a used the primary antibodies anti-GFP ab290 (Abcam) at 1: 5000 dilution, and anti-RAB-A2a^25^ (at 1:1000 dilution, respectively. Alkaline-phosphatase-coupled goat anti-rabbit secondary antibody (Sigma Aldrich) and Western Blue stabilised substrate (Promega) were used to visualise blotted proteins. For immunoblots against endogenous Kin-12F, crude protein extracts from 7 day old *Arabidopsis* seedlings were used. Extracts were probed on western blots with anti-kin-12F polyclonal antibody at 1:1000 concentration (or equivalent concentration of pre-immune serum or immune-depleted serum). Blotted proteins were then visualised using goat anti-rabbit HRP secondary antibody (Thermo Fischer Scientific) at dilution 1: 900 and ECL Western HRP substrate (Merck Millipore).

### Generation of anti-Kin-12F antibody

The polyclonal antibody was raised against the peptide sequence APPQNPNIHNPRNQSV, which is specific to the N-terminus of Kin-12F. This peptide was synthesised as [C]-APPQNPNIHNPRNQSV-amide, and subsequently used for immunisation of two rabbits and ELISA tests (Biosynth Laboratories). Antisera were derived from 50mL harvest bleeds. The crude antisera was purified by affinity chromatography on Thiopropyl Sepharose 6B coupled with the peptide antigen (Biosynth Laboratories). Bound antibodies were eluted in Glycine buffer (100mM, pH 2.5).

### Immunofluorescence

Root were fixed and stained with antibodies and DAPI following previously published procedure^51^. The immuno-depleted anti-Kin12F was prepared by incubating the serum diluted 1:500 with the recombinant Kin-12F antigen at final concentration 0.2 μg/ml for 30 minutes at room temperature. For staining, the anti-Kin-12F antibody was diluted in 1x PBS supplemented with 0.5% (w/v) BSA at 1:100 dilution. Roots were then incubated with this antibody solution or immunodepleted solution overnight at 4°C, washed three times for 1 hour each with 1xPBS at room temperature, and subsequently incubated with rat monoclonal anti-tubulin YL-1/2 antibody (Sigma-Aldrich) at 1:100 dilution. The anti-tubulin was washed out as above, and the roots were incubated overnight at 4°C with a mixture of secondary anti-rabbit conjugated with CY3 and anti-rat conjugated with CY2 (Jackson ImmunoResearch) each diluted to final concentration 1:500. The secondary antibodies were washed out as above except that the second wash contained 50 ng/ml of DAPI to stain the DNA.

### Gliding Assays

The motor domains of Kin-12A, -12B and -12F were cloned from the *Arabidopsis* cDNA and integrated into a vector carrying sfGFP and 6×His sequences (pET-23c backbone). Expression of the proteins was induced in SoluBL21 E. coli with 0.2 mM IPTG for 20 h at 18°C. Harvested cells were lysed using the Advanced Digital Sonifier D450 (Branson) in lysis buffer (25 mM MOPS [pH 7.0], 250 mM KCl, 2 mM MgCl2, 1 mM EGTA, 20 mM imidazole, 0.1 mM ATP) supplemented with 5 mM β-mercaptoethanol, 500 U benzonase and protease inhibitors (1 mM PMSF and peptide inhibitor cocktail: 5 mg/mL aprotinin, 5 mg/mL chymostatin, 5 mg/mL leupeptin, 5 mg/mL pepstatin A). After centrifugation, the clarified lysate was incubated with nickel-NTA coated beads for 1.5 h at 4 °C. Following three washes with the lysis buffer, proteins were eluted using 500 µl elution buffer (25 mM MOPS [pH 7.0], 75 mM KCl, 2 mM MgCl2, 1 mM EGTA, 200 mM imidazole, 0.2 mM ATP) supplemented with 5 mM β-mercaptoethanol. The elution was aliquoted after the addition of 20% (w/v) sucrose.

Microtubule polymerisation was performed by preparing a tubulin mixture. This contained 70% pig tubulin and 30% Alexa Fluor 568-labeled pig tubulin in a total concentration of 100 µM with 0.5mM GMPCPP, which was prepared in MRB80 (80 mM Pipes-KOH, pH6.8, 1mM EGTA, 4 mM MgCl2). The tubulin mixture was incubated for 35 min at 37°C to induce polymerisation. For the gliding assay with purified kinesins, the method followed that previously described^39^, and 1×Standard Assay Buffer (SAB: 25 mM MOPS [pH 7.0], 75 mM KCl, 2 mM MgCl2, 1 mM EGTA) was utilised. Briefly, 7 µL of the purified recombinant protein was introduced into a flow chamber and incubated at room temperature for 2 min in the dark. Subsequently, a 10 µL reaction mixture (1×SAB, 0.1% methylcellulose, 50 mM glucose, 0.5 µg/µL κ-casein, GMPCPP-stabilised microtubule seeds, oxygen scavenger system, 1 mM ATP) was introduced into the flow chamber, and it was sealed using candle wax. TIRF imaging of Alexa Fluor-568 labelled GMPCPP-stabilised microtubule seeds was performed every 3 s for 10 min at 23–25 °C.

### rBiFC

For rBiFC assays, *Agrobacterium* cultures were grown overnight, washed and then resuspended in a solution of 100mM MgCl_2_, 10µM acetosyringone. Cultures were then infiltrated into the abaxial side of *Nictotiana benthamiana* leaves at an O.D 600 of 0.1 for single infiltrations and of 0.2 for co-infiltrations (0.1 for each construct). Plants were imaged 48 hours after infiltration.

### Microscopy and image analysis

Confocal microscopy was performed using a Zeiss 980 CLSM with Airyscan 2 using a C-Apochromat 40x/1.20 W Corr M27 objective. Experiments monitoring phragmoplast expansion over extended time periods used imaging chambers as described before^52^. Imaging of GFP, YFP, RFP, and FM4-64 was as previously described^25^. Brightfield images of GUS-stained primary roots were acquired using a Zeiss Axio Imager.M2 upright microscope. Brightfield images of whole *Arabidopsis* seedlings in Figures 3, S3 and 5 were acquired using a Leica MZ12 microscope with Axiocam. Image processing and analyses (sectioning, reslicing, maximum intensity projections, image assembly, and quantification) were performed using Fiji^53^.

To quantify phragmoplast expansion rates, 3D confocal stacks acquired at 5-minute intervals were resliced with an output spacing of 0.6µm from the top and then from the left. If necessary, images were straightened to align with the vertical plane between top and left reslicing. Oval sections were then drawn around the phragmoplast(s) at successive time intervals until an oval could no longer reliably be drawn around the phragmoplast (because of fusion of one or more sections with the cell periphery). From these oval sections, the area and Feret diameters were calculated. Phragmoplast expansion rates were calculated as the change in Feret diameter between the start and end of the observation period, relative to the time elapsed.

For the quantification of fluorescence intensity at cell plates in longitudinal direction, CLSM stacks or single midplane sections of primary roots co-expressing fluorescently tagged proteins of interest were collected at Nyquist resolution (voxel size 99.5nm x 99.5nm x 550nm). Midplane sections of meristematic cells were imported into Fiji, and cell plates were manually traced using a freehand line. A plot profile with a width 7 pixels was generated and fluorescence intensity was measured along the profile. Average signal intensity was calculated for 1µm wide intervals along cell plates, which were grouped by diameter into diameter into short (<9µm), medium (9-13µm) and long (>13µm). Based on our morphological quantifications of phragmoplasts, these categories correspond to disk stage (short), ring stage (long) and transitioning phragmoplasts (medium). Average intensity +/-SD was plotted using the ggplot2 function in R^54^. Fluorescence intensity orthogonal to cell plates was quantified from the same type of midplane sections described above, plot profiles were generated from straight lines positioned orthogonal to the cell plate with a width of 30 pixels. These were positioned at the centre of the cell plate for disk stage divisions, and at one of the leading edges for ring-stage divisions. Average signal intensity was calculated for 0.25µm intervals, and average intensity +/-SD was plotted using the ggplot2 function in R^54^.

For quantification of phragmoplast morphology, CLSM stacks of primary roots expressing fluorescently RFP:TUB6 were collected at Nyquist resolution (voxel size 99.5nm x 99.5nm x 550nm). Midplane sections of meristematic cells were generated in Fiji, and a freehand line was traced around the phragmoplast. Minimum and maximum Feret diameters were calculated for each phragmoplast in Fiji, which correspond to phragmoplast width (minimum Feret diameter) and diameter (maximum Feret diameter). Double box plots and violin plots for disk and ring-stage phragmoplasts were generated using the ggplot2 function in R^54^.

For quantification of rBIFC signals, CLSM stacks of *Nicotiana benthamiana* leaves co-expressing spit-YFP and mRFP1 were imported into Fiji, and maximum intensity projections of epidermal cells were made. Cells were manually segmented from maximum intensity projections using the polygon tool in Fiji^53^, and mean intensity of YFP and mRFP1 were measured. To count the number of punctae per cell, the area outside each cell wall was cleared in Fiji, and particles were detected using the “Analyse Particles” tool with an intensity threshold of 85, a size filter of 0.20-50.00, and a circularity of 0.00-2.00. Punctae per area were calculated based on the particle number divided by cell surface area, and both mean YFO signal and punctae per area were normalised by nRFP1 intensity to account for differences in expression level in each cell.

### Statistical data analysis and plotting

Two-way ANOVA (analysis of variance) was performed in R using the aov function from the stats package^55^. Tukey’s test was performed in R using the TukeyHSD function from the stats package, Student’s t-test were performed in R using the t.test function from the stats package. Box-, Ribbon-, and Violin-plots were generated in R using the ggplot2 function^54^. In box plots, the median is displayed as a line, lower and upper hinges correspond to the 25th and 75th percentiles, the lower and upper whiskers extend from the hinge to the smallest or largest value no further than 1.5 * IQR from the hinge. Data beyond the end of the whiskers were plotted individually. Violin plots show the same information as box plots, with the addition of the kernel probability density of the data at different values. Ribbon blots show the data mean +/- standard deviation (shaded areas).

## Primer table

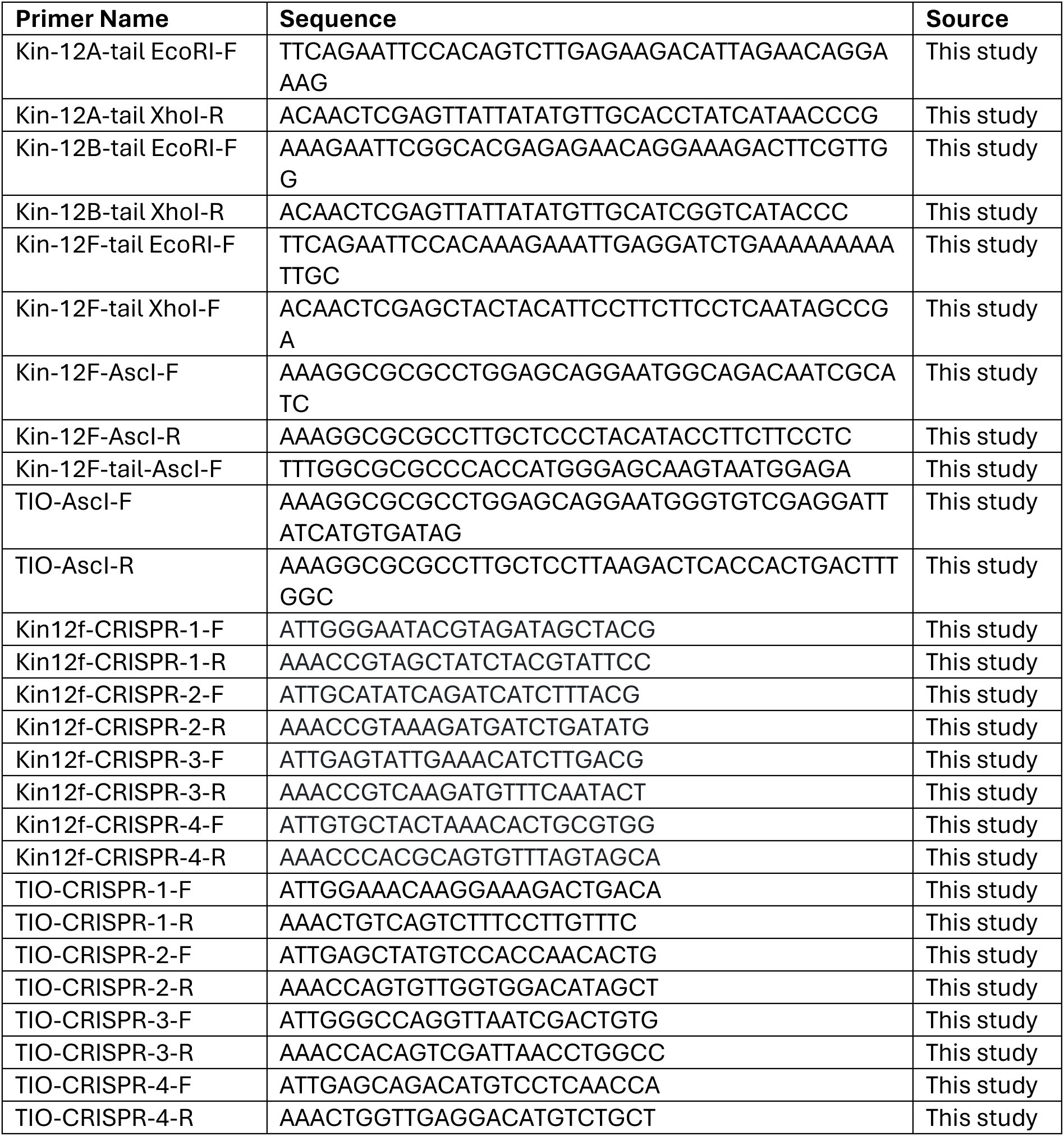
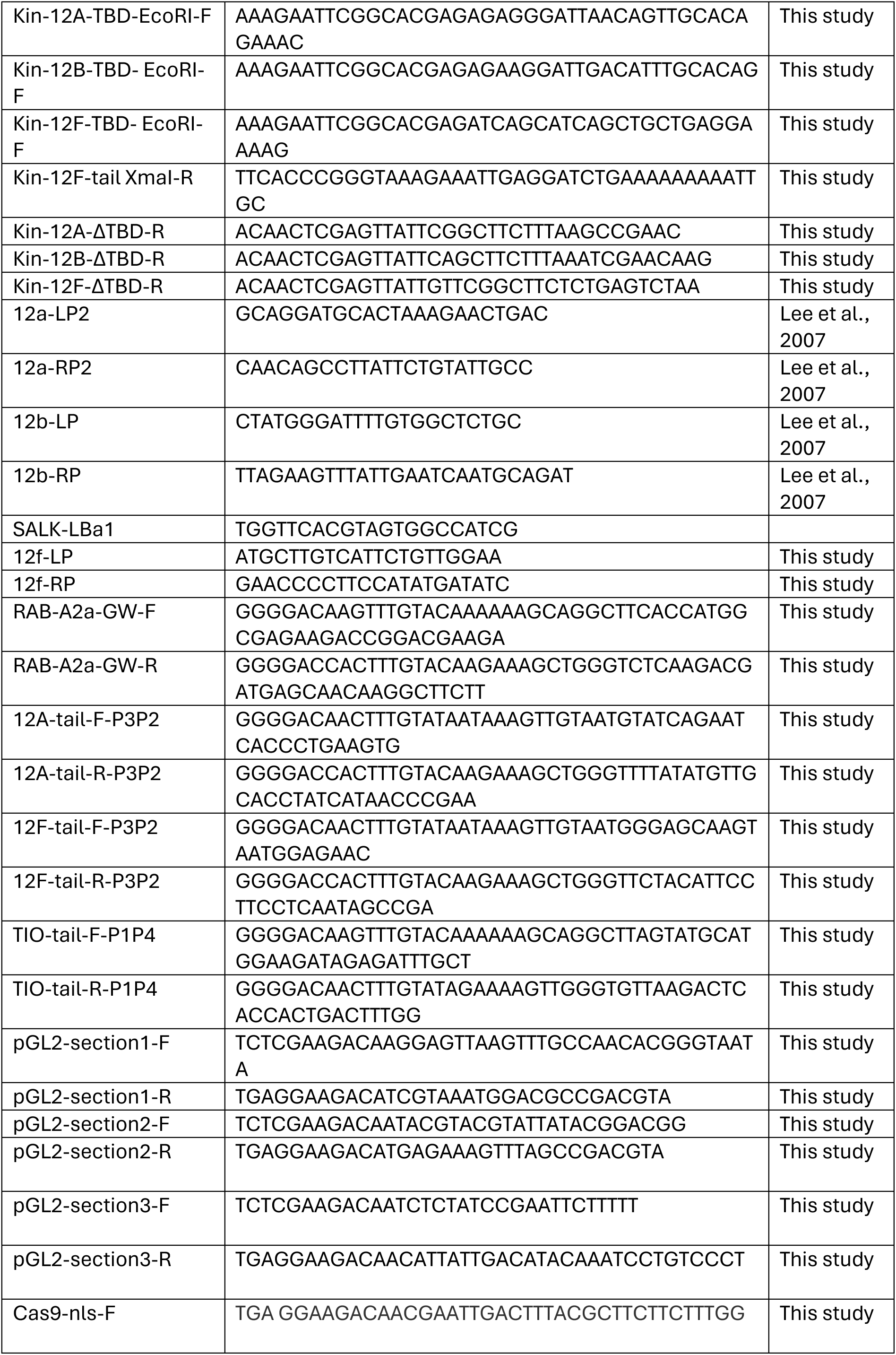
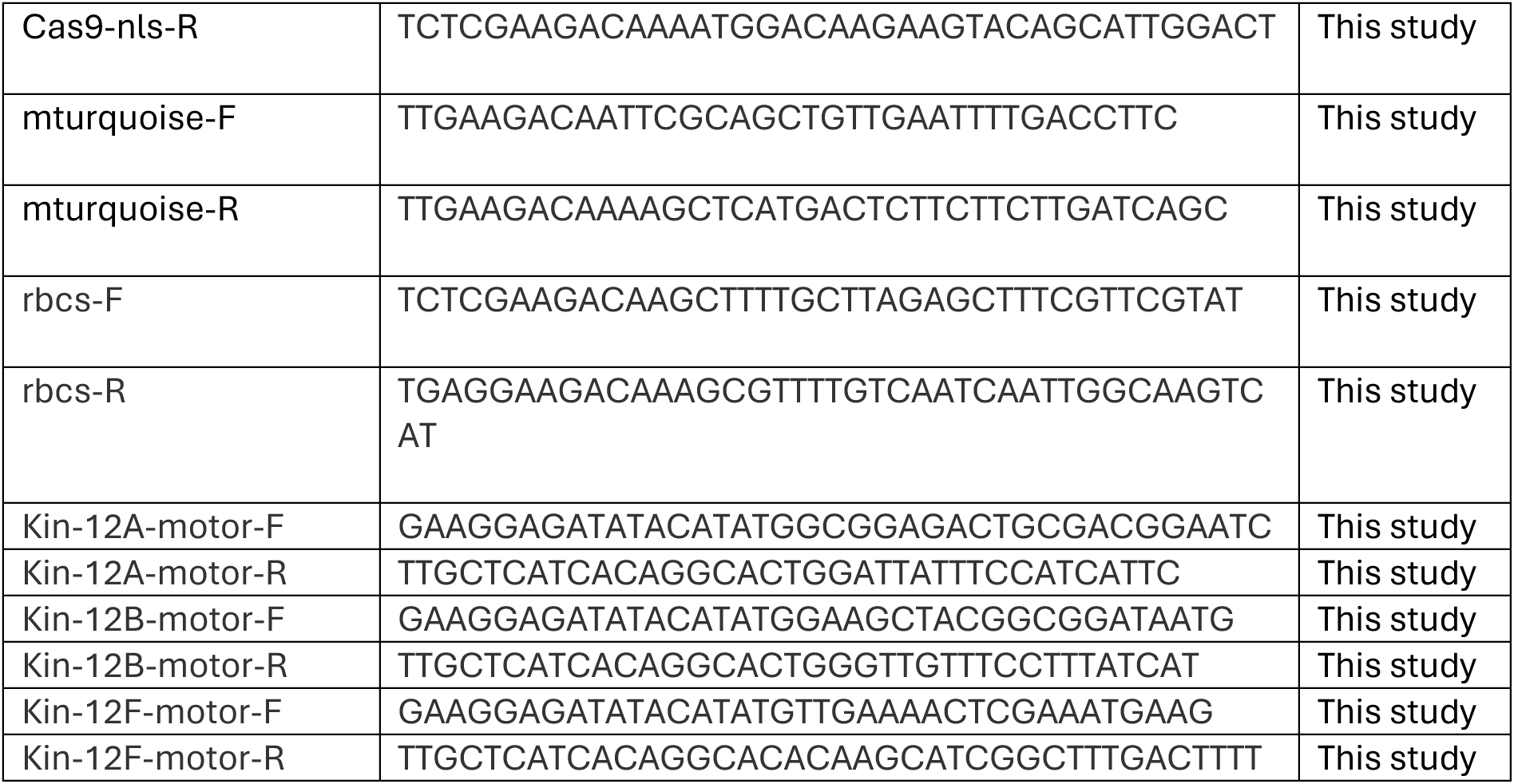

## Acknowledgements

We are grateful to Marie-Cécile Caillaud for published materials and useful discussions, as well as to Sung Aeong Oh and Yoan Coudert for published materials. We are also indebted to Gohta Goshima for useful discussions and facilitating interactions. We further thank Olivier Hamant and Marie-Cécile Caillaud useful feedback on this manuscript. We acknowledge funding from the Biotechnology and Biological Sciences Research Council in the form of a studentship (no. 1810136) to L.E. and grant no. BB/P01979X/1 to C.K. and I.M., as well as funding from the European Research Council (ERC-2020-Stg 948514—EDGE-CAM) to C.K. The work of M.H. and V.Z. on plant cytokinesis is supported by the Czech Science Foundation project 23-05564S.

## Author contributions

Conceptualisation: L.E., I.M., and C.K. Formal analysis: L.E., and C.K. Funding acquisition: L.E., I.M., and C.K. Investigation: L.E., M.K., M.Y., M.H., J.E., and A.S. Methodology: L.E., I.M., M.Y., F.R., Y.J., and C.K. Project administration: I.M. and C.K. Resources: F.R., and N.I. Supervision: I.M., V.Z., P.J.H., Y.J, and C.K. Writing – original draft: L.E. and C.K. Writing – editing: L.E., P.J.H., M.H., V.Z., Y.J., A.S., and C.K.

## Image licenses

Graphics created using biorender.com in this paper were done so by Liam Elliott through a Postdoc Plan subscription, agreement number DY274PGY8B.

## Supplementary Figures

**Supplementary Figure 1:**
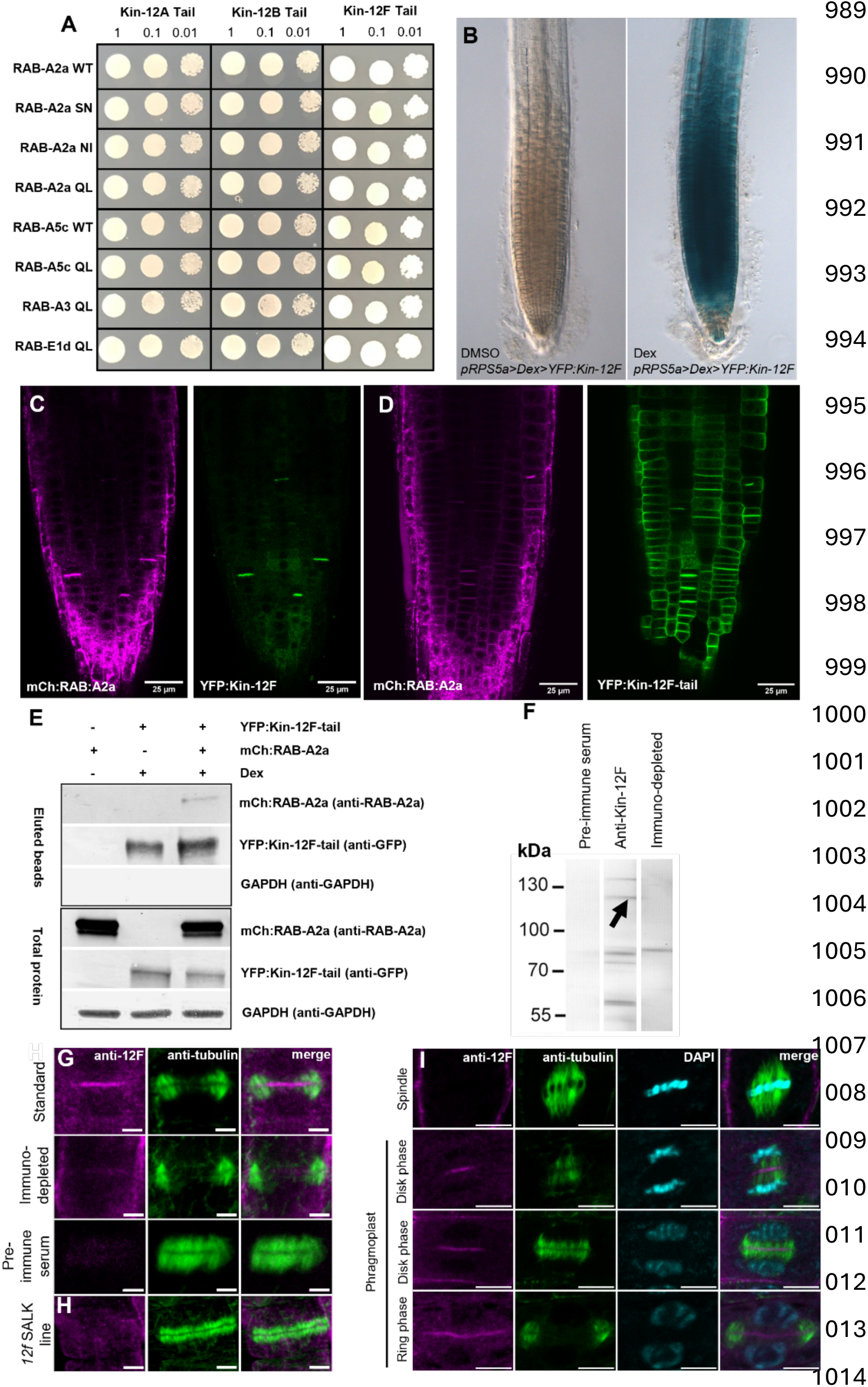
RAB-A2a interacts with class II kinesin-12 members *in vitro* and *in planta*. (A) Pairwise Y2H tests between Kin-12 tail regions and Rab-A GTPase variants on SD-Leu-Trp. Related to Figure 1B. (B) Brightfield images of GUS-stained DEX>> YFP: Kin-12F primary roots after 16hr treatment with either 5µm Dex or DMSO. (C&D) CLSM section of primary roots co-expressing mCh: RAB-A2a and DEX>>YFP: Kin-12F (C) or DEX>>YFP: Kin-12F-tail (D). (E) Immunoblots analysed with anti-YFP, anti-RAB-A2a and anti-GAPDH showing co-immunoprecipitation between YFP: Kin-12F-tail and mCh: RAB-A2a in lines co-expressing DEX>>YFP: Kin-12F-tail and mCh: RAB-A2a, in presence (+) or absence (-) of Dex to induce expression of YFP: Kin-12F-tail. (F) Immunoblots analyses of *Arabidopsis* seedling protein extract with pre-immune serum, anti-kin12F, and immune-depleted serum. Arrowhead indicates band corresponding to Kin-12F of ∼125kDa. (G) CLSM sections of endogenous Kin-12F immunolocalization in primary root meristematic cells using anti-kin12F and anti-tubulin in wild-type plants with standard, immune-depleted and pre-immune conditions. (H) CLSM section of endogenous Kinesin-12F immunolocalization using anti-kin12F and anti-tubulin in a *kin-12f* SALK insertion line. (I) CLSM sections of endogenous Kin-12F immunolocalization in primary root meristematic cells using anti-kin12F and anti-tubulin in wild-type plants counterstained with DAPI. Scale bars 5µm (G, H) and 10µm (I).

**Supplementary Figure 2:**
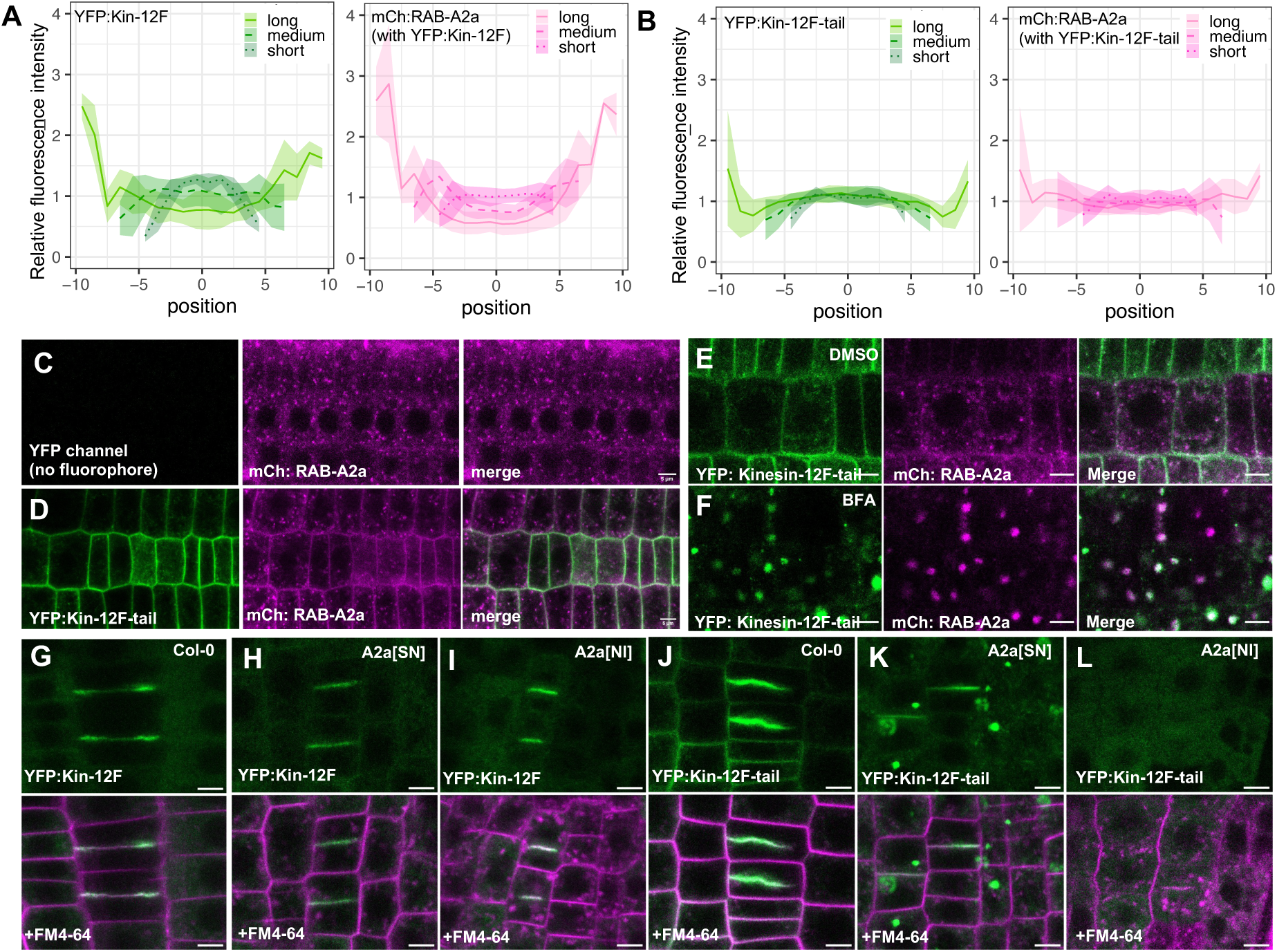
RAB-A2a and Kin-12F mutually determine each other’s localisation patterns. **(A,B)** Fluorescence intensity of YFP:Kin-12F or YFP:Kin-12F-tail and co-expressed mCh:RAB-A2a along cell plates as those shown in Figure 2A,B. Cell plates were grouped by diameter into short (<9µm), medium (9-13µm) and long (>13µm). Lines are mean values, shaded areas are +/- 1SD. N = 4 (YFP:Kin-12F-tail short), 7 (YFP:Kin-12F long), 10 (YFP:Kin-12F short), 11 (YFP:Kin-12F-tail long), 14 (YFP:Kin-12F medium), 16 (YFP:Kin-12F-tail medium). Note this data was used to create the Fluorescence intensity ratio plots in Figure 2C. **(C,D)** CLSM sections of primary root epidermal meristematic cells expressing mCh: RAB-A2a without (C) and with (D) DEX>>YFP: Kin-12F-tail co-expression. **(E,F)** CLSM sections of primary root epidermal meristematic cells co-expressing DEX>>YFP:Kin-12F-tail and mCh:RAB-A2a upon 30mins treatment with 50 µM BFA (F) or equivalent DMSO concentration (E). **(G-I)** CLSM section of primary root meristematic epidermal cells expressing DEX>>YFP:Kin-12F alone (G) or with DEX>>RAB-A2a[SN] (H) or DEX>>RAB-A2a[NI] (I) for 16hrs, counter-stained with FM4-64. **(J-L)** CLSM section of primary root meristematic epidermal cells expressing DEX>>YFP:Kin-12F-tail alone (J) or with DEX>>RAB-A2a[SN] (K) or DEX>>RAB-A2a[NI] (L) for 16hrs, counter-stained with FM4-64. Scale bars, 5µm.

**Supplementary Figure 3:**
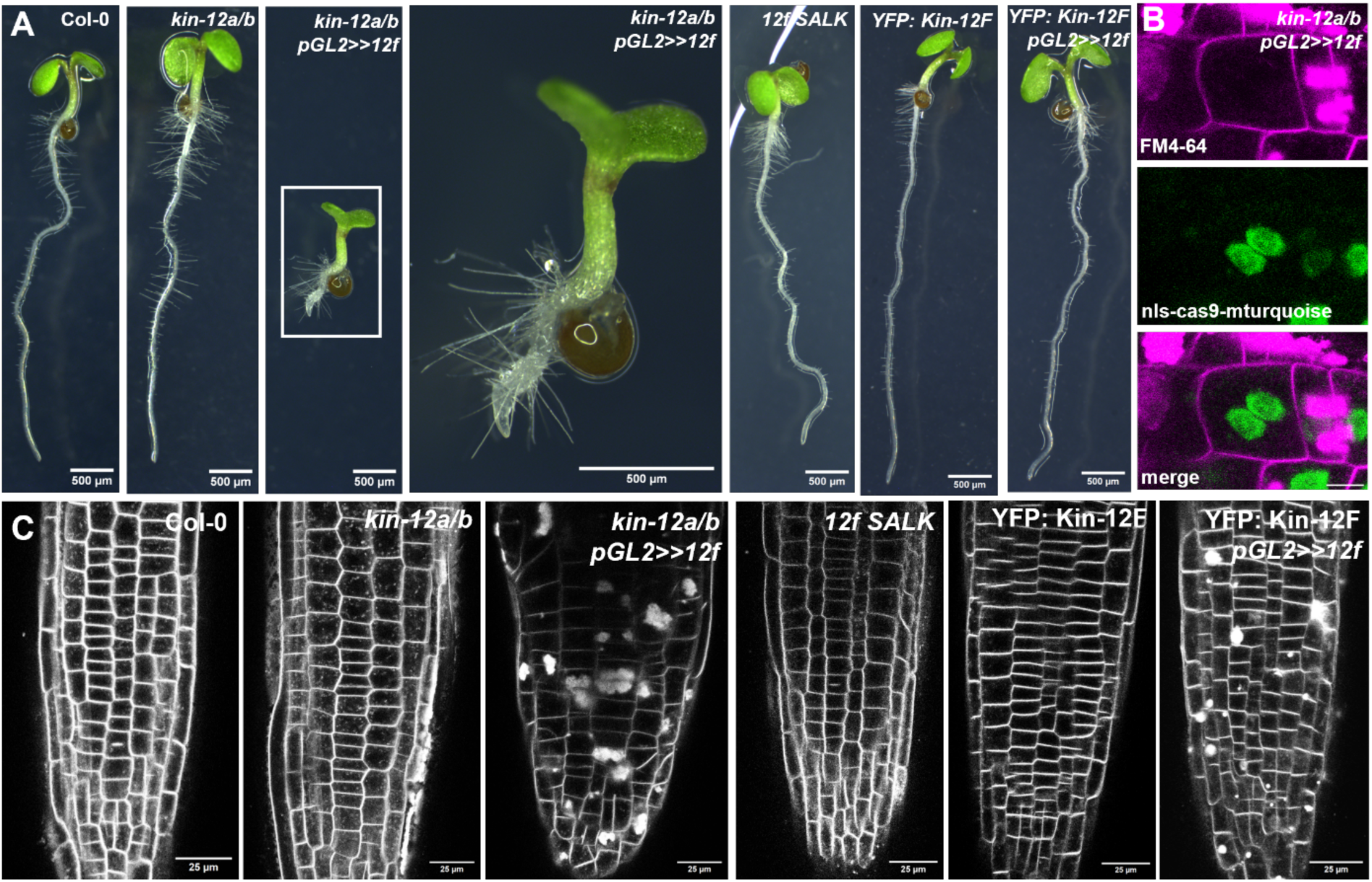
Class II Kin-12 proteins required for somatic cytokinesis and contribute to midzone membrane targeting. **(A)** Brightfield images of 5 day old seedlings of Col-0, *kin12a kin12b, kin12a kin12b pGL2>>12f, 12f SALK, YFP: Kin-12F,* and *YFP: Kin-12F pGL2>>12f* backgrounds. **(B)** CLSM sections of *kin12a kin12b pGL2>>12f* binucleate primary root epidermal meristematic cell. Scale bar, 10µm. **(C)** CLSM sections of primary roots of Col-0, *kin12a kin12b, kin12a kin12b pGL2>>12f, 12f SALK, YFP: Kin-12F,* and *YFP: Kin-12F pGL2>>12f* backgrounds counterstained with FM4-64.

**Supplementary Figure 4:**
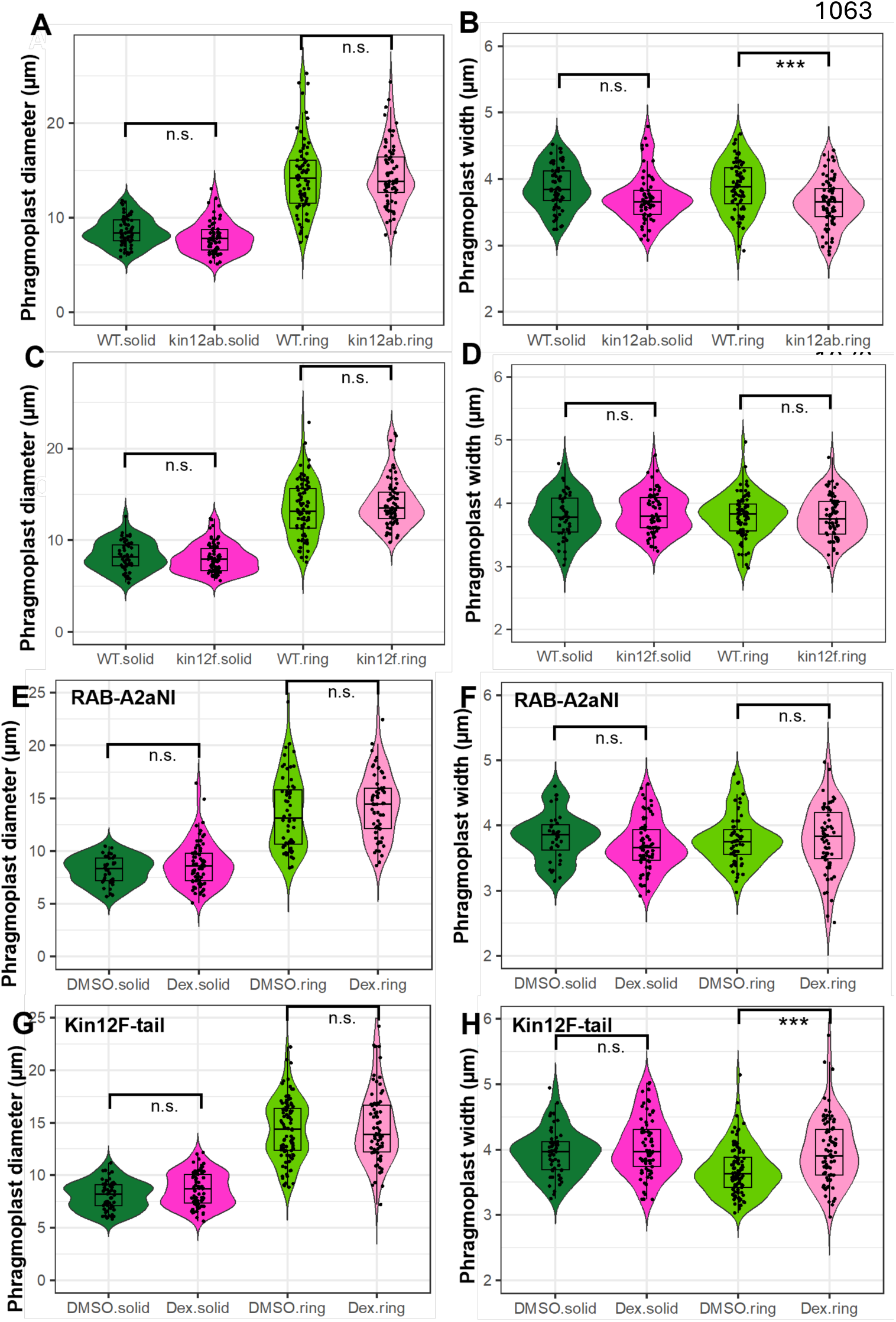
Class II Kin-12 proteins and RAB-A2a are required for phragmoplast expansion. **(A,B)** Violin plots of phragmoplast diameter (A) and width (B) in wild-type and *kin12a/b* backgrounds expressing RFP:TUB6 during disk and ring phase. N = 66 (Col-0 disk), 67 (*kin-12a/b* disk), 73 (Col-0 ring), and 79 (*kin-12a/b* ring). There is a small but statistically significant difference in phragmoplast width between wild-type and *kin12a/b* during ring stage, otherwise phragmoplast morphology is indistinguishable (two-way ANOVA and post-hoc Tukey test, n.s. = p≥0.05; ***=p<0.001). Note data used for violin plots is identical to that used in boxplots in Figure 4B. **(C,D)** Violin plots of phragmoplast diameter (C) and width (D) in wild-type and *kin12f* backgrounds expressing RFP:TUB6 during disk and ring phase. N = 49 (Col-0 disk), 68 (*kin-12f* ring), 75 (*kin-12f* disk), and 83 (Col-0 ring). Phragmoplast morphology is indistinguishable between between wild-type and *kin12f* (two-way ANOVA and post-hoc Tukey test, n.s. = p≥0.05). Note data used for violin plots is identical to that used in boxplots in Figure 4G. **(E,F)** Violin plots of phragmoplast diameter (E) and width (F) during disk and ring phase in presence and absence of DEX>>RAB-A2aNI expression. N = 31 (DMSO disk), 43 (DMSO ring), 50 (Dex ring), and 57 (Dex disk). Phragmoplast morphology is indistinguishable between between DMSO and Dex-treated plants (two-way ANOVA and post-hoc Tukey test, n.s. = p≥0.05). Note data used for violin plots is identical to that used in boxplots in Figure 5B. **(G,H)** Violin plots of phragmoplast diameter (G) and width (H) during disk and ring phase in in presence and absence of DEX>>YFP: Kin-12F-tail expression. N = 57 (DMSO disk), 73 (Dex disk), 78 (Dex ring), and 97 (DMSO ring). Phragmoplast morphology is indistinguishable between between DMSO and Dex-treated plants apart from phragmoplast width at ring stage, which is increased in the presence of Dex (two-way ANOVA and post-hoc Tukey test, n.s. = p≥0.05; ***=p<0.001). Note data used for violin plots is identical to that used in boxplots in Figure 5G.

**Supplementary Figure 5:**
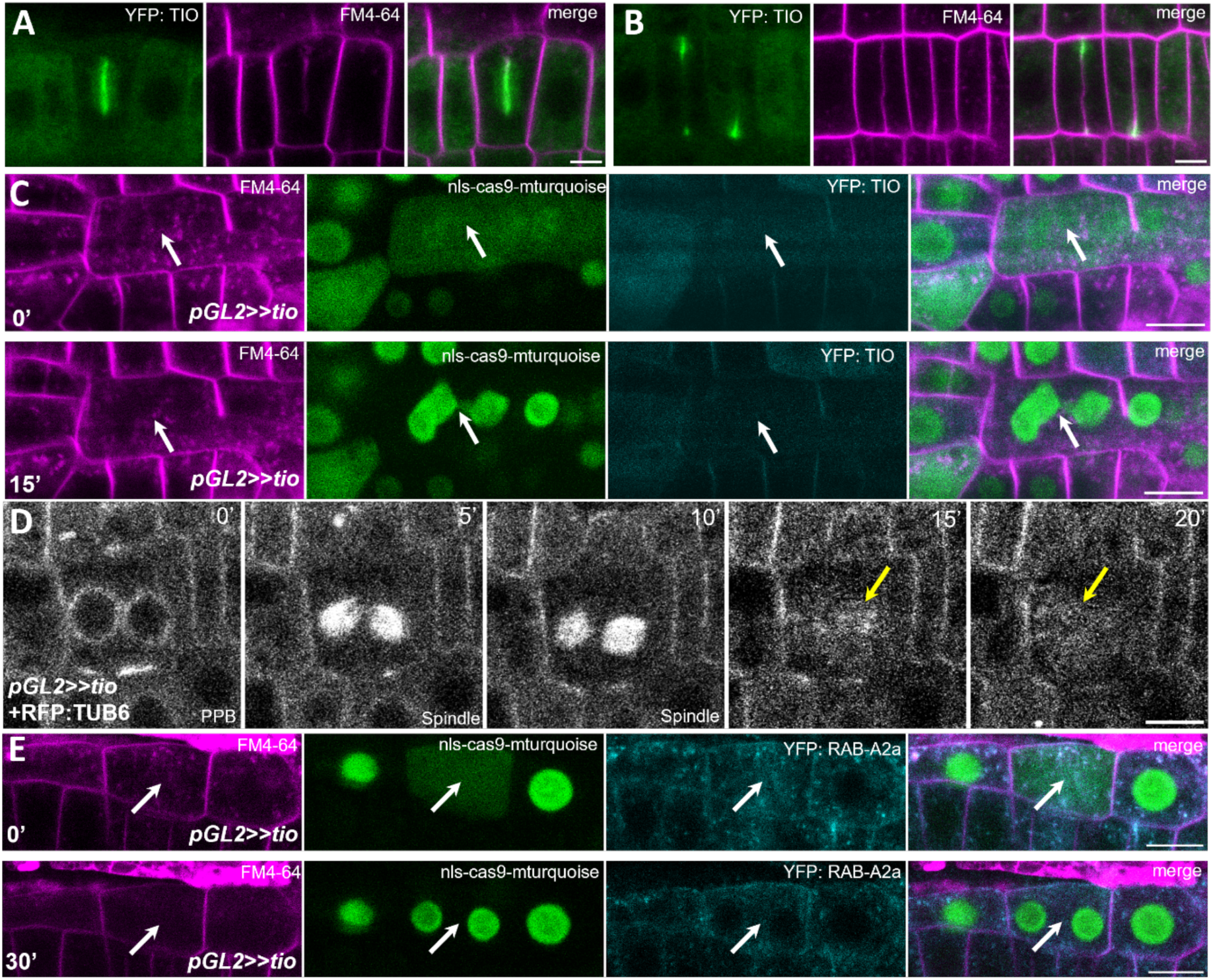
TIO is required for somatic cytokinesis during which it plays a role in phragmoplast formation. **(A,B)** CLSM sections of primary root epidermal meristematic cells expressing DEX>>YFP: TIO counterstained with FM4-64 during early-mid (A) and late (B) cytokinesis. **(C)** CLSM sections of primary root epidermal meristematic cells co-expressing *pGL2::tio::cas9-nls-mturquoise* (*pGL2>>tio)* and DEX>>YFP: TIO before (0 mins) and after (15 mins) nuclear reformation during cell division. White arrow indicates midzone position between future nuclei. **(D)** Sequential CLSM sections of primary root epidermal meristematic cells co-expressing *pGL2>>tio* and RFP: TUB6. Yellow arrow indicates amorphous mass of microtubules at cell centre after spindle dissolution. **(E)** CLSM sections of primary root epidermal meristematic cells co-expressing *pGL2>>tio* and YFP: RAB-A2a before (0 mins) and after (30 mins) nuclear reformation during cell division. White arrow indicates midzone position between future nuclei. Scale bars, 5µm.

**Supplementary Figure 6:**
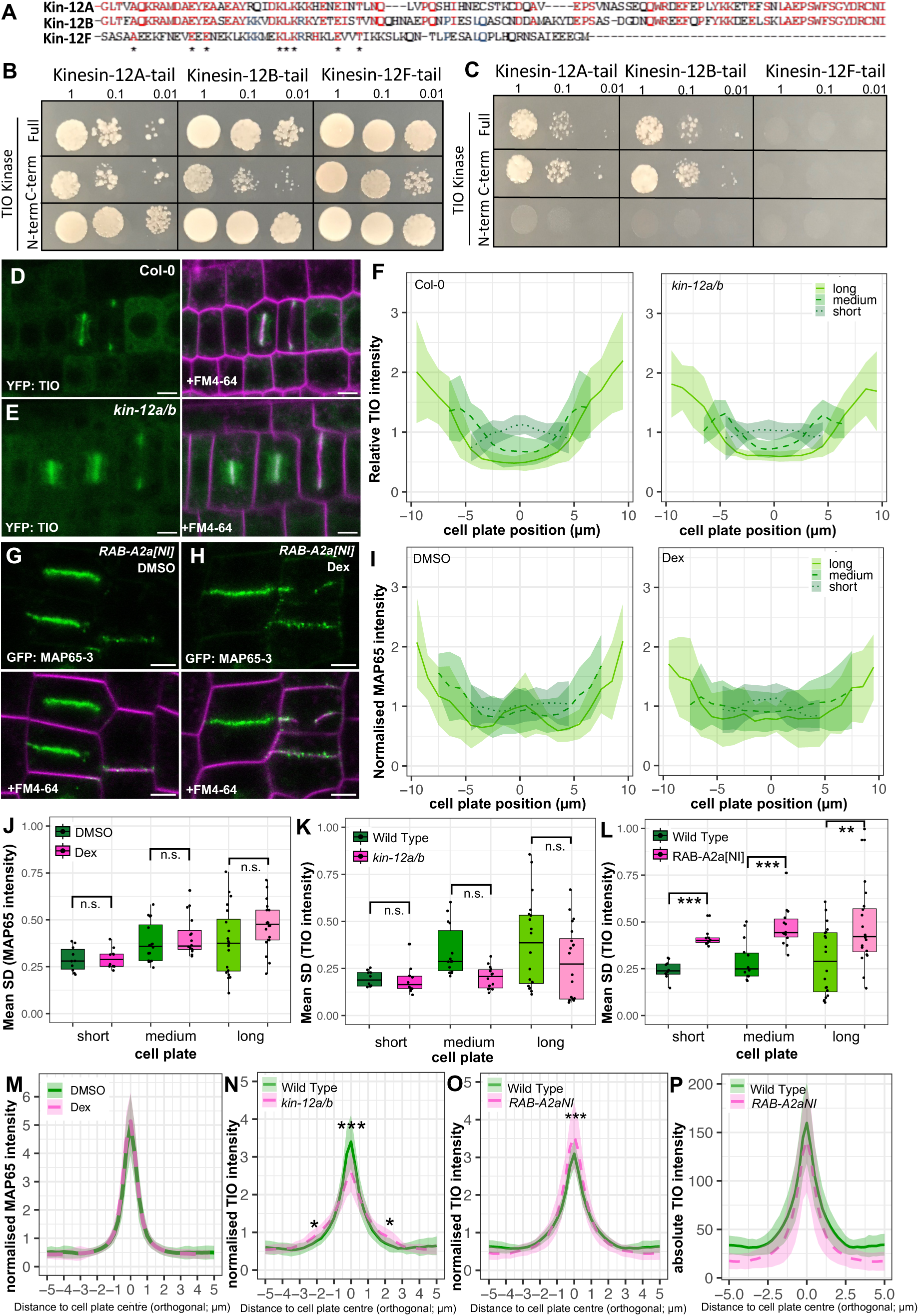
TIO does not interact with Kin-12F, and depends on RAB-A2a and Kin-12A/-12B for its proper localisation. **(A)** Amino acid sequence alignments of conserved residues in known TIO interaction domain of Kin-12A/B compared to Kin-12F. Red indicates residues conserved between Kin-12A/B, blue indicates residues conserved between Kin-12B/F, asterisks indicate residues conserved between all three kinesins. **(B,C)** Pairwise Y2H tests between Kin-12 tail regions and TIO kinase variants on SD-Leu-Trp (A) and SD-Leu-Trp-Ade-His (B). **(D,E)** CLSM sections of primary root epidermal cells expressing YFP:TIO in Col-0 wild type (D) or *kin-12a/b* (E) backgrounds counter-stained with FM4-64. **(F)** Relative fluorescence intensity of YFP:TIO in along cell plates as those shown in (D,E). Cell plates were grouped by diameter into short (<9µm), medium (9-13µm) and long (>13µm). Lines are mean values, shaded areas are +/- 1SD. N = 12 (*kin-12a/b* medium), 19 (Col-0 medium), 23 (Col-0 long), 26 (*kin-12a/b* long), 28 (Col-0 short), and 35 (*kin-12a/b* short). **(G-H)** CLSM sections of primary root epidermal cells expressing GFP: MAP65-3 in absence (G) or presence (H) of DEX>>RAB-A2aNI expression counter-stained with FM4-64. **(I)** Relative fluorescence intensity of GFP: MAP65-3 along cell plates as those shown in (G,H). Cell plates were grouped by diameter into short (<9µm), medium (9-13µm) and long (>13µm). Lines are mean values, shaded areas are +/- 1SD. N = 10 (DMSO long), 13 (Dex long), 30 (DMSO medium), 42 (DMSO short), 55 (Dex short), and 60 (Dex medium). **(J, K)** Mean SD of GFP: MAP65-3 (J) or YFP:TIO (K) fluorescence along cell plates as those in Figure S6D,E,G,H. There is no significant difference in SD of GFP: MAP65-3 distribution (J) in the presence of RAB-A2a[NI] (two-way ANOVA and post-hoc Tukey test, n.s. = p≥0.05), nor in TIO: YFP between wild-type and *kin-12a/b* (K). Scale bar, 5µm. **(L)** Mean SD of YFP: TIO along cell plates as shown in Figure G&H in presence and absence of RAB-A2a[NI] (two-way ANOVA and post-hoc Tukey test, ** = p<0.01; *** = p<0.001). **(M-O)** Mean normalised intensity of GFP: MAP65-3 (M) and YFP: TIO (N, O) in axis perpendicular to cell plate in presence or absence of RAB-A2a[NI] (M, O) or wild-type v.s. *kin-12a/b* backgrounds (N). (P) Mean absolute intensity of YFP: TIO in axis perpendicular to cell plate in presence or absence of RAB-A2a[NI]. Note that while mean normalised YFP: TIO intensity is significantly enhanced at the midzone in the presence of RAB-A2a[NI], mean absolute YFP: TIO intensity is not, demonstrating that the difference in relative signal at the midzone is driven by reduced background signal in the presence of RAB-A2a[NI]. Scale bars, 5µm.

**Supplementary Figure 7:**
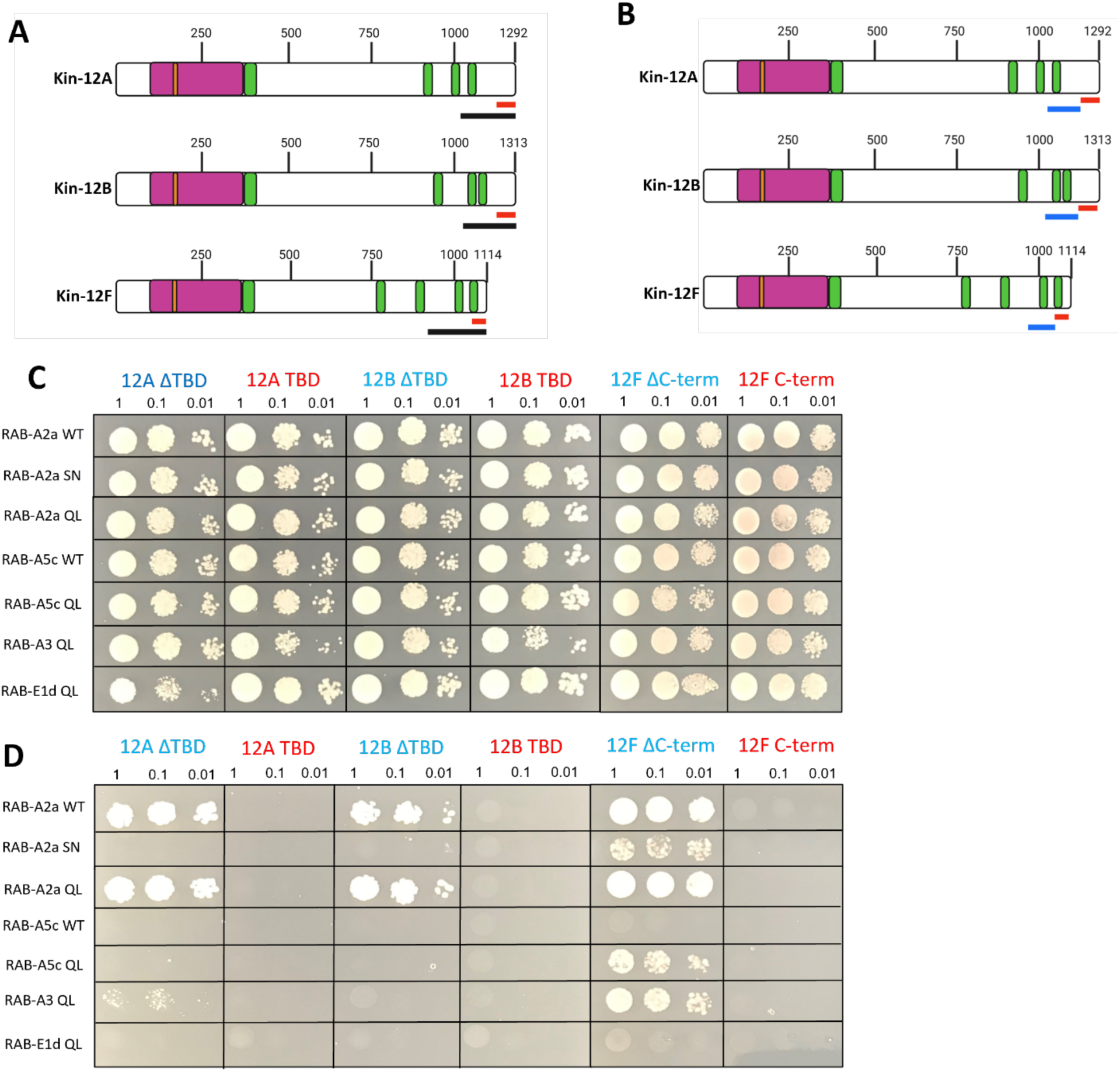
The binding domains of RAB-A2a and TIO on Kin-12A/-12B are adjacent but do not overlap. **(A)** Schematic depiction of Class II Kin-12s structure and binding domains of RAB-A2a and TIO. Black lines: clone region isolated as interactors of RAB-A2a in initial Y2H screen. Red lines: known TIO binding domains on Kin-12A/B and C-terminus of limited homology on Kin-12F. **(B)** Schematic depiction of Class II Kin-12 members structure with refined binding domains of RAB-A2a and TIO based on Y2H tests in (C,D). Blue lines: refined interaction domains of RAB-A2a (12A/B ΔTBD and 12F ΔC-term). Red lines: known TIO binding domains on Kin-12A/B and region of limited homology on Kin-12F (12A/B TBD and 12F C-term). **(C,D)** Pairwise Y2H tests between Kin-12 tail region truncations and Rab-A GTPase variants on SD-Leu-Trp (C) and SD-Leu-Trp-Ade-His (D). Graphics created with biorender.com

**Supplementary Figure 8:**
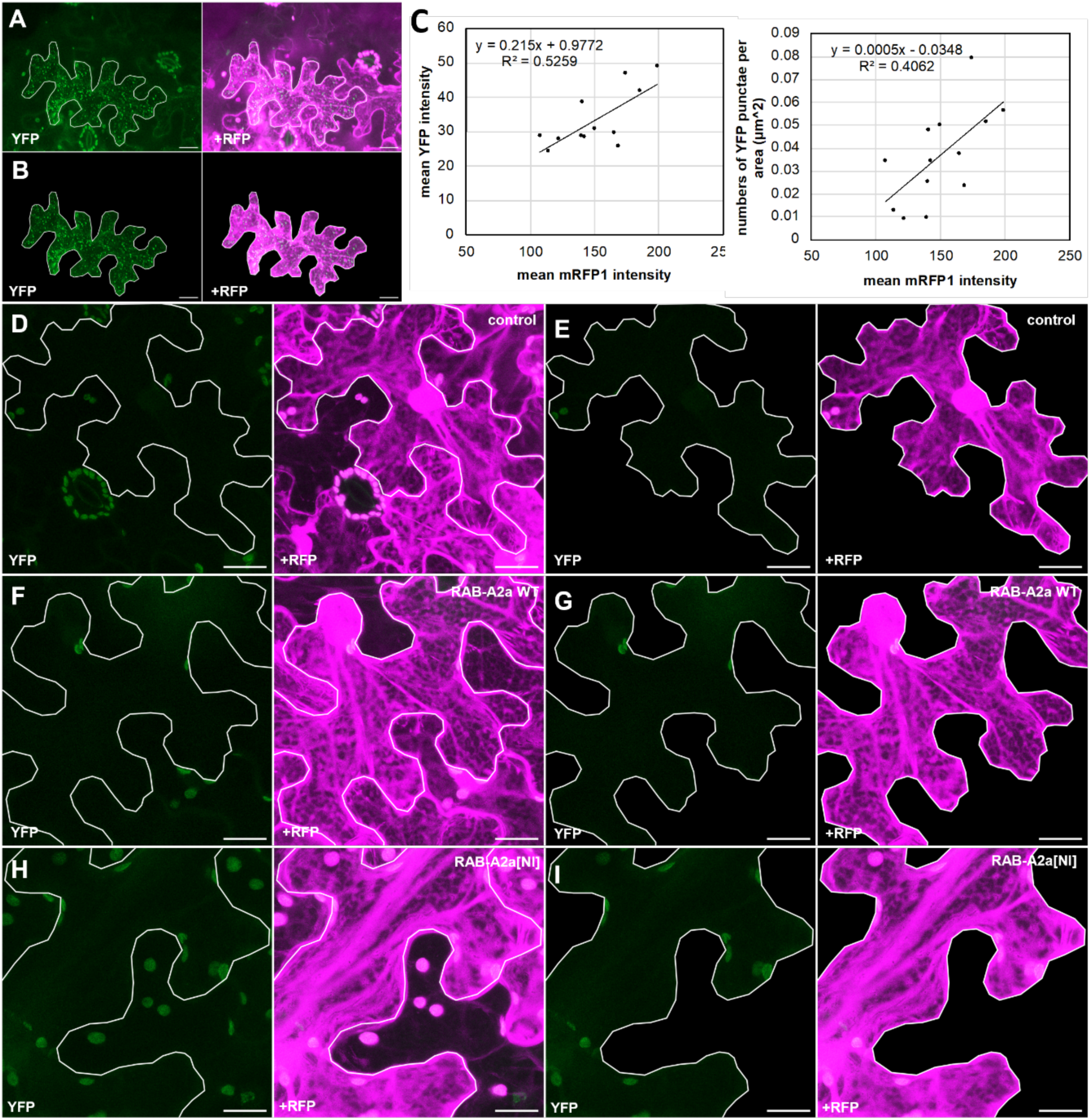
Kin-12F does not interact with TIO in rBiFC assays. **(A)** CLSM maximum intensity projections of *Nicotiana benthamiana* epidermal leaf cells transiently expressing nYFP:TIO and cYFP:Kin-12A and mRFP1 from a ratiometric BiFC system before (A) and after (B) segmentation of cells. **(C)** Correlation of mean YFP intensity or mean YFP punctae per area with mean mRFP1 intensity from cells as those shown in (A,B). **(D-I)** CLSM maximum intensity projections of *N. benthamiana* epidermal leaf cells transiently expressing nYFP:TIO and cYFP:Kin-12F and mRFP1 from a ratiometric BiFC system on its own (D,E) or alongside RAB-A2aWT (F,G) or RAB-A2aNI (H,I) before (D,F,H) or after (E,G,I) cell segmentation. Note no YFP signal was detected, but chloroplast autofluorescence is visible. Scale bars, 20µm

